# Nuclear compression-mediated DNA damage drives ATR-dependent Lamin expression and mouse ESC differentiation

**DOI:** 10.1101/2024.03.15.585216

**Authors:** Tanusri Roy, Swetlana Ghosh, Niyati Piplani, Lakshmi Kavitha Sthanam, Niharika Tiwary, Sayak Dhar, W. Chingmei Wangsa Konyak, Santosh Surendra Panigrahi, Priya Singh, Divya Tej Sowpati, Sreelaja Nair, Sushil Kumar, P. Chandra Shekar, Shamik Sen

## Abstract

Embryonic stem cells (ESCs) which are susceptible to DNA damage depend on a robust and highly efficient DNA damage response (DDR) mechanism for their survival. However, the implications of physical force-mediated DNA damage on ESC fate remains unclear. We show that stiffness-dependent spreading of mouse ESCs (mESCs) induces DNA damage through nuclear compression, with DNA damage causing differentiation through early induction of Lamin A/C expression. Interestingly, differentiation is associated with rescue of DNA damage and activation of the DDR factor ATR. While ATR is typically known to play roles in DDR pathway, its role during stiffness-mediated nuclear compression and mESC differentiation is unknown. Nuclear enrichment of activated ATR on stiff substrates and reduction of Lamin A/C expression upon ATR inhibition suggests that mESC differentiation is driven by nuclear compression-mediated DNA damage and involves ATR-dependent modulation of Lamin A/C.

## Introduction

Embryonic stem cells (ESCs) isolated from the inner cell mass of a late blastocyst of a developing embryo ^1^, serve as an important tool for studying pluripotency and development apart from stem cell-based therapies and genetic engineering ^2^. Due to their high rate of proliferation and a shorter G1 phase ^3,4^, ESCs are particularly susceptible to DNA damage, and depend on a robust and highly efficient DNA damage response (DDR) mechanism to maintain their self-renewability and pluripotency ^5^. ATM (Ataxia telangiectasia mutated) and ATR (ATM and Rad3-related) are key players entailed during DDR which are not only activated by DNA damage signals but also implicated in oxidative stresses ^6,7^, and metabolic processes ^8,9^. While ATM senses DNA double stranded breaks (DSBs), ATR senses single stranded DNA that occurs due to replication stalling and DSB resection^10^. Single stranded DNA have been shown to activate ATR at DNA damage sites, which in turn activates Chk1^11^. However, DSBs with single stranded overhangs interfere with ATM activation and induce an ATM to ATR switch at these damage sites^12^. In addition, ATR also serves as a critical regulator during embryonic development, disruption of which can cause chromosomal fragmentation and lethality in early embryos^13^ and premature age-related phenotypes in adult mice^14^.

The dynamic stem cell microenvironment or the stem cell niche regulates stem cell fate through chemical (secretory) as well as physical cues necessary for survival and proliferation ^15–20^. Force transmission to the nucleus through the linker of nucleoskeleton and cytoskeleton (LINC) complex modulates expression of mechanosensitive genes through chromatin reorganization ^21–23^ that directs stem cell differentiation ^24^ through extracellular matrix (ECM) stiffness ^19^ and topography ^20^. Lamin A/C – a critical component of the LINC complex, not only plays an important role in maintenance of genome integrity ^25–28^ but also controls and contributes to nuclear mechanotransduction and stem cell fate determination ^29–32^. Lamin A/C have been reported to maintain the naïve pluripotency of ESCs by being a critical regulator of chromatin organization while preventing aberrant cell fate decisions^33^. The nuclear lamina, majorly composed of lamins is subjected to dynamic assembly and disassembly by phosphorylation of Lamin A/C during mitosis ^34^. Phosphorylation of Lamin A/C can also be induced by oxidative stress^35,36^, heat shock response^37^ as well as in response to low ECM stiffness^38^. Lamin A/C phosphorylation at Ser22 has been shown to cause its degradation^38^ and nuclear softening which is crucial for protease independent migration of cancer cells ^39^.

Extreme nuclear deformation has been shown to cause nuclear membrane rupture leading to genomic instability ^40–42^. Beyond their canonical DNA repair functions in somatic cells, recent studies have documented ATM/ATR activation in cells subjected to mechanical stresses ^43–46^. While alterations in ATR can cause nuclear envelope breakdown and aberrant chromatin organization ^43,44^, loss in ATM triggers nuclear deformation and reduced Lamin A levels thus regulating nuclear stiffness and mechanics ^47^. Given the association between ECM stiffness and chromosomal instability in mouse ESCs (mESCs) ^48^, in this study, we have probed the inter-relationships between ECM stiffness, DNA damage, DNA repair and mESC fate. By culturing mESCs on substrates of increasing stiffness, we show stiffness-dependent mechanoadaptation induces DNA damage through increased nuclear compression. This DNA damage in turn triggers stiffness dependent induction of Lamin A/C expression. While ATR plays a widespread role, we identify ATR as a regulator of Lamin A/C expression in mediating initiation of differentiation by obstructing Lamin A/C phosphorylation. Collectively, our study establishes stiffness-induced nuclear compression that mediates early ESC differentiation and ATR-dependent modulation of Lamin A/C expression.

## Materials & Methods

### Cell Culture & Reagents

E14 Tg2a mouse embryonic stem cells (mESCs) were cultured on 0.1% porcine gelatin (Sigma) coated dishes in presence of KO-DMEM (Gibco) and supplemented with 10% KO-Serum replacement (Gibco), 1X Glutamax (Gibco), 1X non-essential amino acids (NEAA) (Gibco) and 1X antibiotic-antimycotic (Gibco). These cells were supplemented with 10ng/ml leukemia inhibitory factor (LIF) and 0.1mM 2-mercaptoethanol (Gibco) to maintain pluripotency. Mouse embryonic fibroblasts (MEFs) were cultured in DMEM (Gibco) supplemented with 10% Fetal Bovine Serum (FBS) of South American origin (Gibco) and 1X antibiotic-antimycotic (Gibco). Both mESCs and MEFs were maintained at 37^0^C and 5% CO_2_ concentration (NUAIRE incubator). For experiments, cells were dislodged with 1X TrypLE Express (Gibco) and seeded on glass or MEFDMs or polyacrylamide hydrogels (PA gels) coated with either 5µg/cm fibronectin (Merck) or 10µg/cm^2^collagen-I (Sigma) without LIF. mESCs were treated with etoposide (ETO) (2µM, Sigma), Blebbistatin (Blebb) (5µM, Sigma), MnCl_2_ (0.5 mM), N-acetyl cysteine (NAC) (5 mM, Sigma), Retinoic acid (RA) (1µM, Sigma), Ascorbic acid (AA) (50µM, Sigma), KU-55933 (10µM, Sigma) and VE-821 (5µM, Sigma) ^49^ for a period of 24 hours.

### Fabrication of Polyacrylamide (PA) gels

PA hydrogels of varying stiffness of 0.6 kPa, 4 kPa and 33 kPa were fabricated by mixing varying ratios of 40% Acrylamide (Bio-Rad) and 2% Bis-Acrylamide (Bio-Rad) with milliQ water as described elsewhere ^50,51^. Gels were polymerized on either 12mm,18mm or 60mm glass coverslips for immunostaining, comet assay and western blotting/flow cytometry experiments respectively. Gels were coated with either 10µg/cm² rat-tail collagen-I (Sigma) or 5µg/cm^2^ human-plasma fibronectin (Merck) overnight in 4°C, post functionalization using Sulfo-SANPAH (Pierce).

### Fabrication of MEF derived matrix (MEFDM)

MEFDMs were generated by seeding WT MEFs (on 1% gelatin coated dishes (Porcine gelatin, Sigma) at a density of 4.6 x 10^5^/cm^2^ as described elsewhere in ^48^.

### Immunostaining and Microscopy

For immunostaining, cells were fixed after 3 hours, 6 hours, 24 hours or 72 hours of culture using 4% PFA (Sigma) in PBS for 15 minutes, permeabilized with 0.1% TritonX-100 (Sigma) in PBS for 10 minutes, and then incubated with one or more primary antibodies overnight at 4°C post blocking with 5% FBS for 1 hr in room temperature. Primary antibodies used for immunostaining were against pMLC2-Ser19 (1:250, Cell Signaling Technologies), γ-H2AX (1:500, Cell signaling Technologies), Lamin A/C (1:400, Cell Signaling Technologies), Lamin B1 (1:500, Abcam), 53BP1 (1:2500, Invitrogen), pATM-Ser1981 (1:500, Invitrogen) and pATR-Thr1989 (1:500, Invitrogen). The following day, cells were washed with PBS and then incubated with one or more secondary antibodies at room temperature for 2 hours. Secondary antibodies used were Alexa-fluor 488 anti-mouse IgG (1:1000, Invitrogen), Alexa-fluor 555 anti-rabbit IgG (1:1000, Invitrogen). Cells were then stained with fluorescently labeled wheat-germ agglutinin (1:1000, Sigma) to stain cell surface and nuclei were labeled with DAPI (1:2000, Sigma). Cells were imaged at 120X magnification using either an inverted fluorescent microscope (Olympus, IX83) or at 63X magnification using Spinning Disc Confocal Microscope (Zeiss, CSU-X1) for greater resolution. Cell spreading area, DNA damage foci, mean intensities, nuclear enrichment and nuclear height were quantified using Fiji ImageJ software while nuclear volumes were quantified using Imaris X64 8.3.1. Stiffness and time-dependent Lamin A/C distributions were generated based by normalizing raw intensity values with the maximum value across all conditions and time-points. The normalized intensities (I*_N_*) on a given stiffness gel (g) were categorized as low (Lo), Moderate (Mo) and High (Hi) based on the following criteria: Lo: I*_N_* ≤ µ*_g_* - σ*_g_*, Mo: µ*_g_* - a*_g_* < I*_N_* ≤ µ*_g_* + σ_*g*_, Hi: I*_N_* > µ*_g_* + σ*_g_*, where µ*_g_* and σ*_g_* correspond to the mean and the standard deviation of normalized intensity observed at the 3 hr time-point. The frequency curves were generated using Origin 2021. The percentage of Lo, Mo and Hi cells were calculated thereafter.

### Comet Assay

To quantify the level of DNA damage in mESCs, alkaline and neutral comet assays were performed according to manufacturer’s protocol (Trevigen). For this, mESCs were cultured on PA gels of varying stiffnesses. Post 24 hours, these cells were isolated from the gels using TrypLE Express (Gibco). 10L cells per ml were suspended in PBS and were mixed with 1% low melting agarose (GeneI) cooled to 37°C at a 1:10 v/v cell solution to agarose ratio. Approximately 50µl of the cell-agarose mixtures were evenly spread and incubated on comet slides (Trevigen) at 37°C for 5-10 minutes, and then transferred to 4°C in dark for 30 minutes until the agarose polymerized. The slides were then incubated overnight in ice cold lysis solution (Trevigen) at 4°C followed by alkaline denaturation for 1 hour to detect both single and double stranded DNA breaks. Electrophoresis was carried out at 330 mA in the presence of ice-cold alkaline electrophoresis solution (pH>13) followed by 70% ethanol fixation for 5 minutes. To exclusively detect double stranded DNA breaks, neutral comet assay was performed by subjecting the cell-agarose mixture comet slides to run in presence of neutral electrophoresis solution (pH = 9) at 1V/cm, after overnight lysis at 4°C (same as alkaline assay). This was followed by a DNA precipitation step for 30 minutes and then fixation in 70% ethanol. Samples were stained with SYBR green dye (1:10,000, Invitrogen) diluted in 1X TE buffer and visualized under the microscope at 10X magnification. Comets were analyzed by feeding tif images into the OpenComet plugin in ImageJ Fiji software. Extent of DNA damage was quantified by comparison of the olive tail moment of mESCs on varying substrate stiffnesses.

### AFM

Cell stiffness was measured using an Atomic Force Microscope (MFP3D, Asylum). Cells were probed with 10 kHz soft, pyramidal silicon nitride probes (Olympus) of nominal stiffness 20 pN/nm. The exact cantilever stiffness was determined using the thermal calibration method. Estimates of cell stiffness were obtained by fitting the first 500 nm of experimental force-indentation curves with Hertz model ^52^. Similarly, the stiffness of gels was estimated by fitting the first 1000 nm of the experimental force-indentation curves as described elsewhere ^51^.

### Western Blotting

mESC protein samples were isolated from mESC pellets using ice-cold RIPA buffer (Sigma) in the presence of freshly added 1% protease and phosphatase inhibitor cocktail (Invitrogen). Cell lysates were cleared by centrifugation at 12,000 rpm for 15 minutes at 4°C. Protein concentration was determined by Lowry method. 60 µg of total protein sample was loaded into each well and resolved by SDS-PAGE on gradient acrylamide-bisacrylamide gels (6% to 15%). Samples were transferred on to 0.2 µm PVDF membranes (Bio-Rad). Membranes were blocked with 5% BSA in TBST for 1 hour at RT and then incubated one or more of the following primary antibodies overnight at 4°C: Lamin A/C (1:1000, CST), pLamin A/C-Ser22 (1:1000, CST), Lamin B1 (1:1000, Abcam), γH2AX (1:1000, CST), H2AX (1:500, CST), pMLC2-Thr18/Ser19 (1:700, CST), MLC2 (1:700, Abcam), Oct3/4 (1:1000, Abcam and CST), Nanog (1:1000, CST), pATM-Ser1981 (1:1000, Invitrogen), pATR-Thr1989 (1:1000, Invitrogen), ATM (1:1000, CST), ATR (1:1000, CST), RAD51 (1:1000, Abcam), pCHK1-Ser345 (1:1000, CST), CHK1 (1:1000, CST), GATA4 (1:1000, Invitrogen), Brachyury (1:1000, Invitrogen), GATA3 (1:1000, Invitrogen), NeuN (1:1000, CST), pRPA2-Ser33 (1:700, Novus Biologicals) and GAPDH (1:1000, CST). The following day, after washing with TBST, membranes were incubated with HRP conjugated secondary antibody (1:10,000, Invitrogen) for 2 hours at RT. Subsequently, membranes were washed again with TBST and developed using ECL detection kits (Advansta), western BLoT Hyper HRP Substrate (Takara) and ultra-sensitive chemiluminescent substrate (Cynagen) for low abundant proteins. Blots were also subjected to stripping using western BLoT stripping buffer (Takara) according to manufacturer’s protocol, followed by reprobing. Images were captured using a chemidoc imaging system (Bio-Rad).

### RNA Sequencing (RNASeq) and heat map generation

Total RNA from bulk cells was isolated using TRIzol (Ambion) according to instructions from the manufacturer. The concentration and purity of samples was checked using the Nanodrop 2000 spectrophotometer. Ribo-Zero Plus stranded library preparation kit (Illumina) was used to generate libraries from a total of 1*µ*g RNA for each sample. Library concentration was estimated using Qubit dsDNA High Sensitivity Assay Kit (Invitrogen). The libraries were sequenced on the Illumina NovaSeq 6000 platform to a depth of 50 million paired-end reads. The quality assessment of sequenced reads was evaluated using FASTQC (v0.12.0) followed by mapping them to the Mus_musculus.GRCm39.cdna reference cDNA transcriptome using Kallisto (v0.45.0) for estimating transcript abundance. Data annotation was carried out using AnnotationHub and genes exhibiting a Counts Per Million (CPM) value of less than 1 in at least three samples were excluded from further analysis. The resulting filtered count matrix was then normalized by Trimmed Mean of M-values (TMM) method using edgeR package, and was subsequently log2-transformed to mitigate data heteroskedasticity. Principal Component Analysis (PCA) was performed using the Scikit-learn library of python and a 3D PCA plot generated. Heatmaps were generated using Count matrix values followed by normalizing the data against +LIF control and then applying z-score normalization across each row to get relative gene expression values. The representative matrices were sorted based on gene expression value of average of 72-hour samples.

### Gene Ontology Analysis

Differential gene expression analysis conducted with the PyDESeq2 package identified upregulated and downregulated genes across stiffness conditions at 72 hours, using +LIF as a reference and applying a log2 fold change threshold of ≥1 and an adjusted p-value of ≤0.01. Following gene identification, gene ontology analysis via the gprofiler2 library was utilized to enrich transcription factor pathways. Subsequently, common transcription factor pathways upregulated across three stiffness conditions were identified. Prevalence of genes associated with endoderm, mesoderm, ectoderm and extraembryonic ectoderm lineage specification within each identified transcription factor pathway was quantified and sorted based on this occurrence frequency. The top 20 pathways were ranked based on their p-values and visually represented in a grouped bar chart format.

### Flow cytometry

For cell-cycle analysis, mESCs were pelleted down using TrypLE Express (Gibco) and fixed using 0.25% PFA for 15 minutes on ice followed by permeabilization with 70% methanol in 4°C for at least 1 hour. Cell pellets were washed with PBS to remove residual methanol and then 50µg/ml Propidium Iodide (Himedia) was added to each condition before acquisition for cell cycle measurements on BD FACS Aria Fusion. Gating was done with respect to pluripotent mESCs using FlowJo software. For apoptosis assay, mESC pellets were incubated with FITC conjugated Annexin V and Propidium Iodide in 1X binding buffer as per manufacturer’s protocol (BD Pharmingen). 10,000 events were acquired per sample. Gating was done with respect to unstained control using FlowJo software.

### Pharmacological inhibition assays using zebrafish embryos

*Tg(cmlc2):mCherry* embryos were raised in embryo medium at 28^°^C till 24 hours post fertilization (hpf). At 24 hpf, embryo medium was replaced with embryo medium containing 10µM KU55933 or 10µM VE821. Embryos were kept in the inhibitors for 24 hours at 28^°^C. The next day, the medium was replaced with fresh embryo medium without inhibitors and embryos were left to grow for another day. Heart phenotypes were quantitated and imaged at 48 hpf (24 hours of exposure to inhibitor) and at 72 hpf (24 hours of exposure to inhibitor, followed by 24 hours of recovery). Live images were obtained using Olympus MVX10 and processed using CellSens and ImageJ softwares.

### Statistical Analysis

For statistical analysis, raw data were processed in Origin 2021. The normality of the data was first checked using the Kolmogorov-Smirnov normality test. Based on these results, either parametric or nonparametric tests were subsequently performed. Statistical significance was assessed using unpaired t-tests for comparing two parametric datasets. One-way or two-way ANOVA tests were performed for parametric datasets with multiple conditions, followed by Tukey’s post-hoc test to compare the means. For multiple non-parametric datasets, Kruskal-Wallis ANOVA followed by Dunn’s post-hoc test while Mann-Whitney test for comparing means between two non-parametric datasets were performed. p < 0.05 being considered as statistically significant.

## Results

### Stiffness-mediated mechanoadaptation is associated with DNA damage in mESCs

Long-term culture of mESCs on stiff gelatin-coated coverslips have been shown to accumulate greater aneuploidy compared to that on mouse embryonic fibroblast (MEF) derived matrices, which are much softer ^48^. To probe the link between stiffness-mediated mechanoadaptation and genomic integrity, we cultured mESCs on collagen (Col) and fibronectin (FN) coated 0.6 kPa, 4 kPa and 33 kPa PA gels that span a physiologically relevant stiffness range (Fig. S 1A). As mESCs exhibited greater spreading and focal adhesion intensities on FN coated gels compared to that on Col coated gels, indicative of a more robust mechanoadaptation response on FN coated gels (Fig. S 1A, B), all subsequent experiments were performed on FN coated substrates up to 72 hrs in culture. While mESCs spread minimally on the softest 0.6 kPa gels, spreading increased substantially on the 4 and 33 kPa gels (Fig. 1Ai, ii).

**Figure 1:**
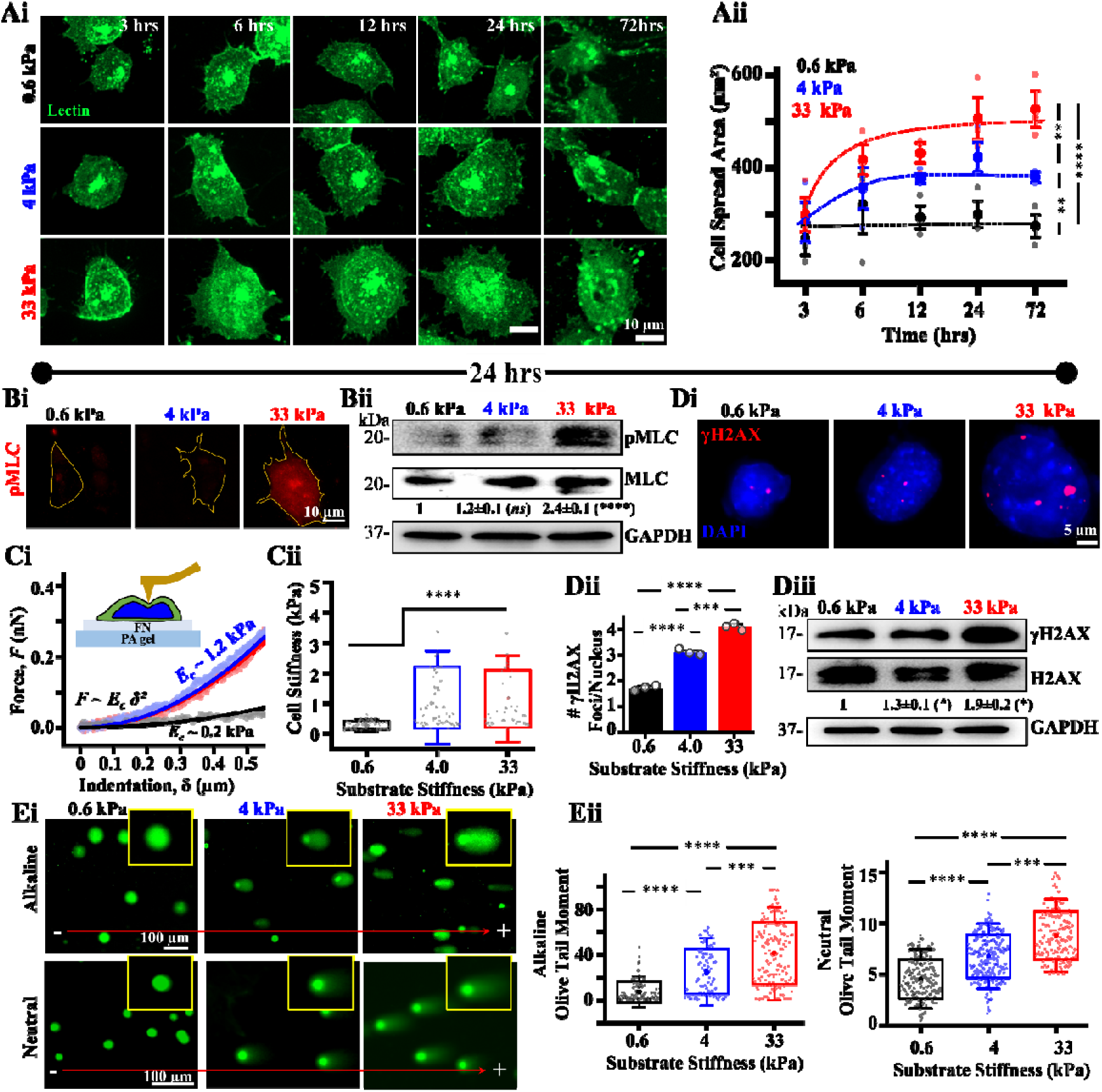
Association of mechanoadaptation and DNA damage in mouse embryonic stem cells (mESCs): (Ai, ii) Representative lectin-stained images of mESCs on fibronectin (FN)-coated polyacrylamide (PA) gels of varying stiffness post 3, 6, 12, 24 hours and 72 hours of seeding and quantification of cell spreading ( cells per condition across *N* = 3 independent experiments). Scale Bar = 10*µm*. Error bars represent SD. Two-way ANOVA with Tukey’s test was used for comparing means (**-value 0.01, ****-value 0.0001). **(Bi, ii)** Assessment of pMLC expression in mESCs cultured on PA gels for 24 hours based on immunofluorescence and western blotting showing mean ±SEM of pMLC/MLC (*N* = 3 independent experiments). Two-way ANOVA with Tukey’s test was used for comparing means with respect to 0.6 kPa conditions (****-value 0.0001). Scale Bar = 10*µm.* **(Ci-iii)** Measurement of mESC cortical stiffness using atomic force microscopy (AFM) after 24 hrs in culture. Cells were probed with a soft pyramidal probe and first 500 nm of the raw force-indentation curves were fitted with Hertz model to estimate cortical stiffness ( cells per condition across *N* = 3 independent experiments). One-way ANOVA with Tukey’s test was used for comparing means (**** p-value <0.0001). **(Di-iii)** Quantification of DNA damage in mESCs cultured on PA gels for 24 hrs. DNA damage was assessed by co-staining cells with DAPI and γH2AX and counting γH2AX foci per nucleus (n > 50 nuclei per condition across *N* = 3 independent experiments) and western blotting of whole cell lysates showing mean ±SEM of γH2AX/H2AX (*N* = 3 independent experiments). Two-way ANOVA with Tukey’s test was used for comparing means with respect to 0.6 kPa conditions (*p-value ≤0.05). Error bars represent SEM. One-way ANOVA with Tukey’s test was used for comparing means (**** p-value <0.0001). **(Ei, ii)** Representative images of alkaline and neutral comet assay of mESCs cultured on PA gels (Insets show single comets) and quantitative analysis of olive tail moment (n > 30 cells per condition across *N* = 3 independent experiments). One-way ANOVA with Tukey’s test was used for comparing means (*p-value <0.05, ** p-value <0.01, ns = non-significant *p*-value>0.05). For all blots, GAPDH served as loading control See also Figure S1 A-C.

Increased spreading on 4 and 33 kPa gels was associated with increased cytoskeletal organization as assessed by active actomyosin contractility (phosphorylated myosin light chain, pMLC) (Fig. 1Bi, ii, Fig. S 1C) and cortical stiffness (Fig. 1 Ci, ii). Interestingly stiffness-mediated mechanoadaptation was associated with stiffness-dependent increase in the extent of DNA damage assessed by counting the number of γH2AX foci in individual cells (Fig. 1 Di, ii) and total protein levels assessed using westerns (Fig. 1 Diii). Since γH2AX marks both single and double strand breaks^53–55^, to determine the relative proportion of single and double strand breaks across different stiffnesses, we performed alkaline comet assay which detects both type of breaks, and neutral comet assay which detects double strand breaks (Fig. 1 Ei, ii). Stiffness-dependent increase in olive tail moments in both the assays suggests the presence of both single and double strand breaks. However, greater stiffness dependent change in olive tail moments in alkaline comet assay compared to neutral comet assay indicates higher proportion of single strand breaks with increase in substrate stiffness. Together, these results establish a direct relationship between stiffness-dependent mechanoadaptation and DNA damage in mESCs.

### DNA damage is induced by mechanoadaptation-mediated nuclear compression

To test if stiffness-induced DNA damage was ROS dependent, DNA damage was assessed by quantifying γH2AX foci in cells cultured in the presence and absence of the ROS scavenger N-acetyl cysteine (NAC). Insensitivity of DNA damage to NAC suggests that mechanoadaptation-mediated DNA damage in mESCs is ROS independent (Fig. 2 Ai, ii). Contractility-mediated nuclear compression has been shown to induce nuclear envelope rupture and DNA damage in cancer cells cultured on rigid glass coverslips ^40^. To check the link between DNA damage and nuclear compression, we captured transverse sections of Lamin B1-stained nuclei of mESCs cultured on different gels and quantified nuclear height. Increased DNA damage on stiffer substrates was associated with increased nuclear compression observed as early as 3 hrs (Fig. 2 Bi, ii). In comparison, nuclear volume increased marginally with time but remained unchanged across different gels (Fig. 2 Biii).

**Figure 2:**
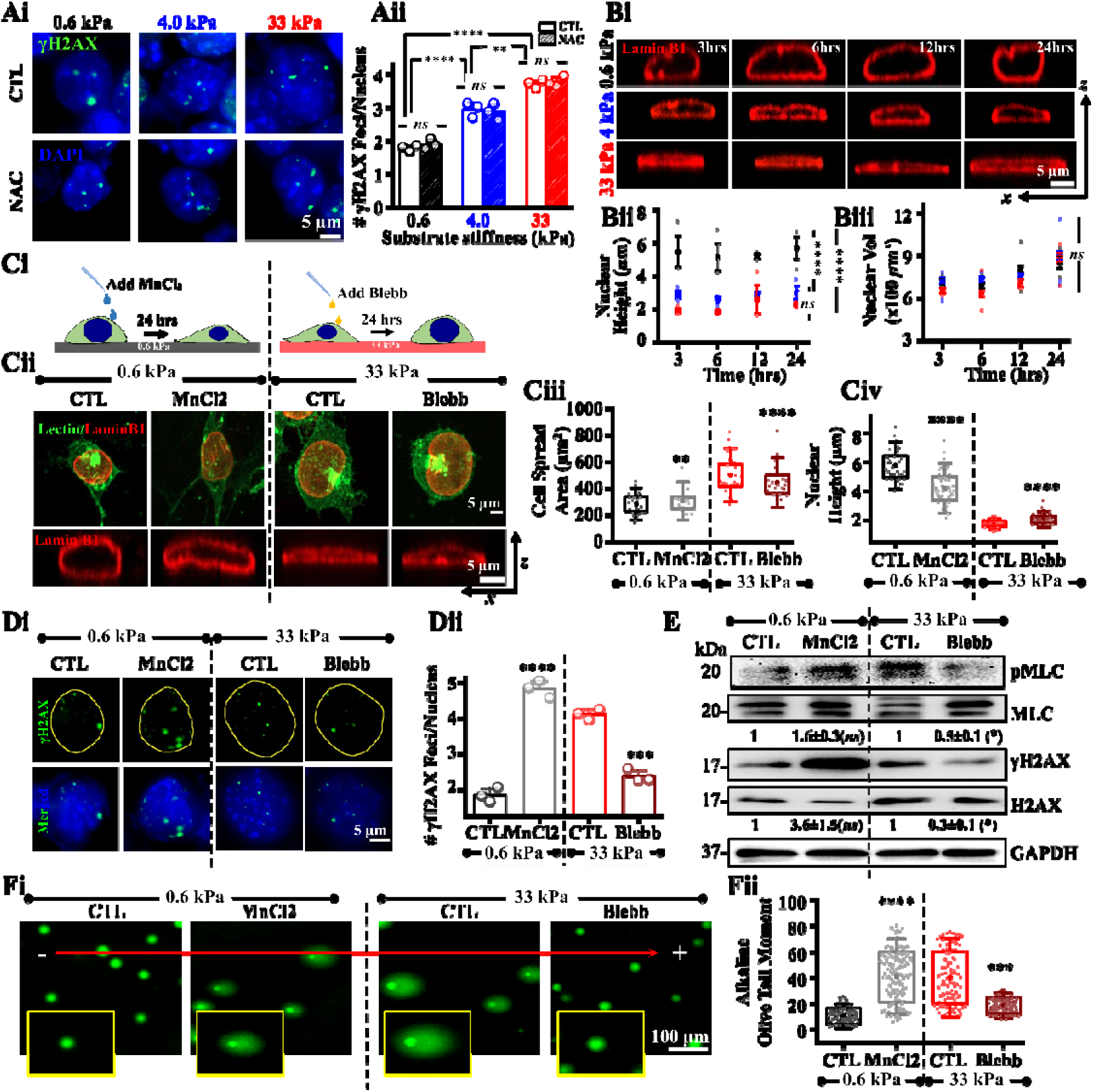
**Mechanoadaptation mediated nuclear compression induces DNA damage in mESCs**: **(Ai, Aii)** Representative images of mESC nuclei co-stained with γH2AX on PA gels of varying stiffnesses after 24 hrs in culture in presence and absence of the ROS inhibitor, N-acetyl cysteine (NAC) and quantification of DNA damage ( cells per condition across *N* = 3 independent experiments). Scale Bar = 5*µm.* Error bars represent SEM. Two-way ANOVA followed by Tukey’s post-hoc test was used to compare means (**-value 0.01, ****-value 0.0001, *ns* = non-significant-value 0.05). **(Bi-Biii)** Representative images showing transverse sections of mESC nuclei immunostained with Lamin B1 on PA gels of varying stiffness post 3, 6, 12 and 24 hours of seeding and quantification of nuclear height and nuclear volume ( cells per condition across *N* = 3 independent experiments). Scale Bar = 5*µm*. Error bars represent SD. Two-way ANOVA with Tukey’s test was used for comparing means (****-value 0.0001, *ns* = non-significant *p*-value 0.05). **(Ci-Civ)** Schematic workflow, representative lectin-stained images and transverse Lamin B1 stained nuclear sections of mESCs on PA gels of 0.6 kPa +/-MnCl_2_ and 33 kPa +/-Blebbistatin and quantification of cell spreading and nuclear height ( cells per condition across *N* = 3 independent experiments). Scale Bar = 5*µm.* Error bars represent SD. Unpaired Student’s t-test was used for comparing means between control and treated samples on each PA gel (**p-value <0.01, **** p-value< 0.0001). **(Di, Dii)** γH2AX and DAPI immunostained images of mESCs on PA gels of 0.6 kPa +/-MnCl_2_ and 33 kPa +/-Blebbistatin. Quantification of γH2AX foci (n> 40 cells per condition across *N* = 3 independent experiments). Scale Bar = 5*µm.* Error bars represent SEM. Unpaired student t-test was used for comparing means between control and treated samples on each PA gel (***p-value <0.001, **** p-value< 0.0001). **(E)** Representative immunoblots and quantification of mean ±SEM of pMLC/MLC and γH2AX/H2AX levels in mESCs cultured on 0.6 kPa gels +/-MnCl_2_ and 33 kPa gels +/-Blebbistatin (*N* = 3 independent experiments). Unpaired Student t-test was used for comparing means with respect to untreated controls of each PA gel (*p-value ≤0.05, *ns* = non-significant p-value>0.05). **(Fi, Fii)** Representative images of alkaline comet assay of mESCs cultured on 0.6 kPa +/-MnCl_2_ and 33 kPa +/-Blebbistatin (Insets show single comets), and quantitative analysis of olive tail moment (n > 30 cells per condition across *N* = 3 independent experiments). Scale Bar = 100 µm. Error bars represent SD. Unpaired Student’s t-test was used for comparing means with respect to untreated controls of each PA gel (***p-value <0.001, **** p-value <0.0001). For all blots, GAPDH served as loading control.

Based on these results, we propose that increased spreading and cytoskeletal organization on stiffer substrates induces DNA damage by nuclear compression. To test this, the extent of nuclear compression was perturbed by modulating cell spreading, and its impact on DNA damage assessed. While cells were treated with MnCl_2_ on 0.6 kPa gels to increase cell spreading via increased integrin association ^56–58^, on 33 kPa gels, cells were treated with the contractility inhibitor blebbistatin (Blebb) (Fig. 2 Ci). On 0.6 kPa gels, MnCl_2_ treatment led to increased spreading (Fig. 2 Cii, iii), decreased nuclear height (Fig. 2 Cii, iv), increased γH2AX foci (Fig. 2 Di, ii) and increase in olive tail moment assayed using alkaline comet (Fig. 2 Fi, ii). Conversely, on the stiff 33 kPa gels, treatment with Blebb led to decreased cell spreading (Fig. 2 Cii, iii), increase in nuclear height (Fig. 2 Cii, iv), reduction in the extent of DNA damage (Fig. 2 Di, ii), and decrease in olive tail moment (Fig. 2 Fi, ii). Western blotting of total cell lysates revealed increased pMLC and γH2AX levels in MnCl_2_-treated cells, and the opposite in Blebb-treated cells (Fig. 2 E). Collectively, these results demonstrate that mechanoadaptation-mediated nuclear compression induces DNA damage in mESCs.

### DNA damage induces loss of pluripotency and Lamin A/C expression

Lamin A/C is an integral part of the nuclear membrane that plays a key role in nuclear mechanics as well as in genome protection. ECM stiffness has been shown to drive differentiation of mesenchymal stem cells (MSCs) with induction of Lamin A/C expression ^19,20,32^ and affect ESC differentiation ^59,60^. However, the importance of stiffness-induced DNA damage in this context remains unclear. Interestingly, increase in Lamin A/C levels on 4 kPa and 33 kPa gels and drop in pluripotency markers Oct3/4 and Nanog suggests that DNA damage may drive early differentiation (Fig. 3 Ai, ii). To test this, experiments were performed wherein DNA damage was induced with low doses of the topoisomerase II inhibitor – etoposide (ETO) and its impact on pluripotency and Lamin A/C expression assessed on the softest 0.6 kPa gels, where mechanoadaptation and DNA damage were lowest (Fig. 3 Bi). Remarkably, ETO treatment led to robust increase in Lamin A/C levels with corresponding loss in Oct3/4 and Nanog expression, suggesting that low doses of DNA damage also induce loss in pluripotency and induction of Lamin A/C expression (Fig. 3 Bii, Biii). To understand the temporal kinetics of Lamin A/C induction triggered by DNA damage, mESCs cultured on 0.6 kPa gels for 6 hrs (to allow for initial attachment) were incubated with 2 µM of ETO, and Lamin A/C levels tracked upto 3 hrs post treatment (Fig. 3 Ci). Both DNA damage and Lamin A/C remained unchanged upto 2 hrs of ETO treatment but increased significantly at the 3 hr time-point (Fig. 3 Cii, iii). The same held true for pluripotency markers, Oct3/4 and Nanog, with significant loss in pluripotency at the 3 hr timepoint.

**Figure 3:**
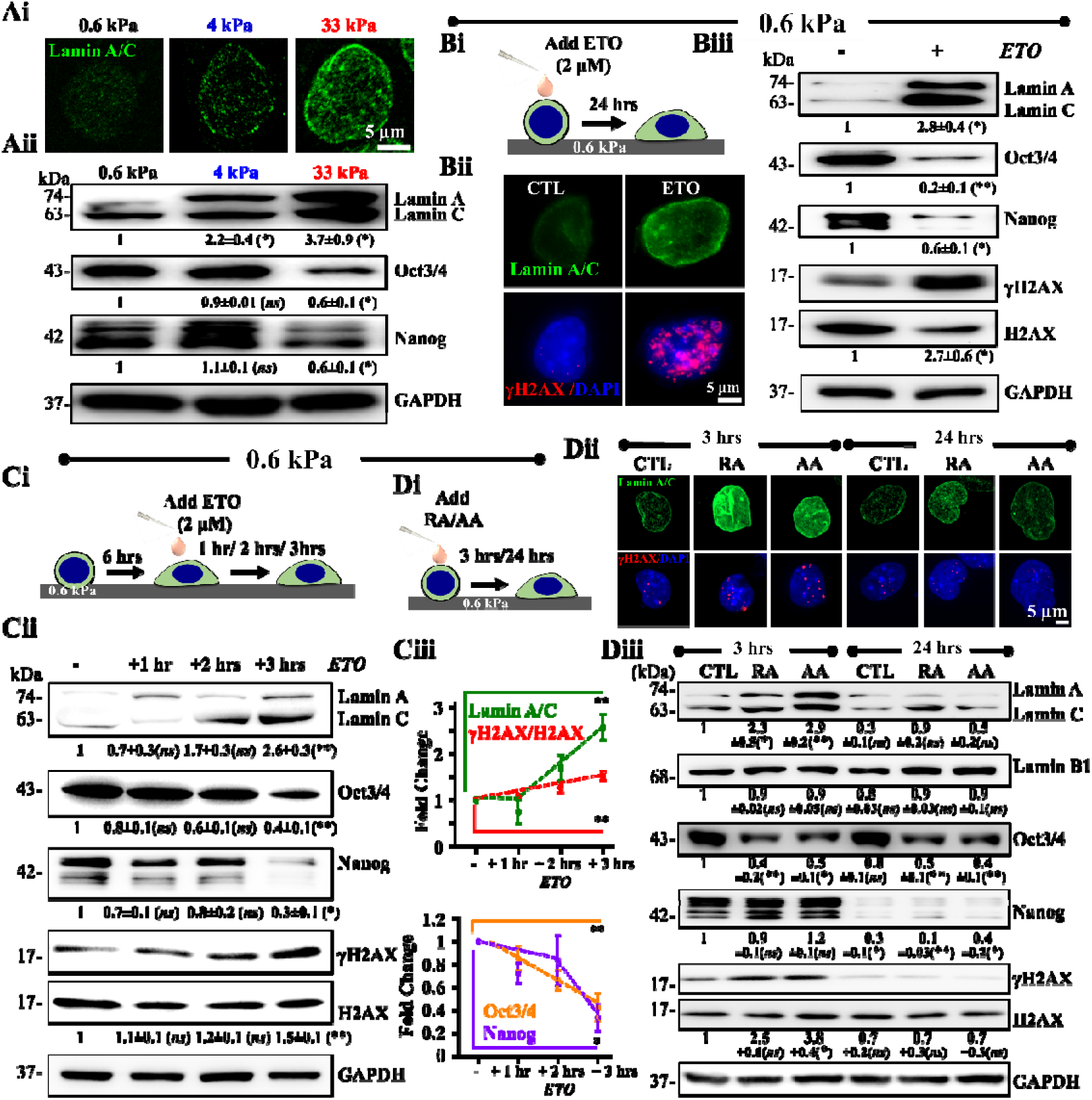
DNA damage induces Lamin A/C expression: (Ai, Aii) Representative images of mESC nuclei immunostained with Lamin A/C on PA gels of varying stiffnesses post 24 hrs of culture. Scale Bar = 5*µm.* Representative immunoblots and quantification of mean ±SEM of Lamin A/C expression and pluripotency (Oct3/4 and Nanog) of mESCs on varying stiffnesses post 24 hrs culture (*N* = 3 independent experiments). Two-way ANOVA with Tukey’s test was used for comparing means with respect to 0.6 kPa conditions (*-value 0.05, *ns* = non-significant-value 0.05). **(Bi, Bii, Biii)** Experimental setup and representative images of Lamin A/C expression and DNA damage in presence of 2µM etoposide (ETO) post 24 hrs treatment on mESCs by immunofluorescence on 0.6 kPa gels. Scale Bar = 5*µm.* Representative immunoblots and quantification of mean ±SEM of Lamin A/C expression, pluripotency levels and γH2AX/H2AX of whole cell mESC lysates in presence and absence of ETO post 24 hrs treatment on 0.6 kPa gels ( independent experiments). Unpaired student’s t-test was used for comparing means with respect to untreated control. (*-value 0.05, **-value 0.01). **(Ci, Cii, Ciii)** Experimental setup for studying temporal correlation between DNA damage and Lamin A/C induction. Cells were allowed to attach onto 0.6 kPa gels for 6 hrs prior to ETO addition, and then cultured for upto 3 hrs. Representative immunoblots and quantification of mean ±SEM showing temporal evolution of Lamin A/C, Oct3/4, Nanog, γH2AX/H2AX (N = 3 independent experiments). One-way ANOVA with Tukey’s test was used for comparing means with respect to untreated controls (*p-value <0.05, **p-value <0.01, *ns* = non-significant *p*-value>0.05). **(Di, Dii, Diii)** Schematic workflow and representative immunofluorescence images of Lamin A/C and γH2AX on mESCs post 3 hrs and 24 hrs of retinoic acid (RA) and ascorbic acid (AA) treatment on 0.6 kPa PA gels. Scale Bar = 5*µm*. Representative immunoblots and quantification of mean ±SEM of Lamin A/C, Lamin B1, Oct3/4, Nanog and γH2AX/H2AX of whole cell mESC lysates post treatment of RA and AA on 0.6 kPa PA gels (N = 3 independent experiments). Two-way ANOVA with Tukey’s test was used for comparing means with respect to 3 hr CTL conditions (*p-value <0.05, **p-value <0.01, *ns* = non-significant *p*-value>0.05). For all blots, GAPDH served as loading control. See also Figure S 1 Di, ii and Figure S 2 A, B.

To assess if chemical factor induced differentiation also involves DNA damage, mESCs cultured on the softest 0.6 kPa gels in the presence of the neuronal differentiation factor retinoic acid (RA) and the osteogenic differentiation factor ascorbic acid (AA) were probed for DNA damage and Lamin A/C levels at 3 and 24 hr time-points (Fig. 3 Di). Both RA and AA treatments led to increased levels of the neuronal marker, β-III tubulin at the 24 hr time-point (Fig. S 1Di, ii). Strikingly, both RA and AA treatment led to increase in γH2AX and Lamin A/C levels, and loss in Oct3/4 levels at the early 3 hr time-point (Fig. 3 Dii, iii). Though Nanog was unchanged at the 3 hr time-point, near complete loss in its expression was observed at 24 hrs indicative of differentiation. Surprisingly, at this timepoint both γH2AX and Lamin A/C levels dropped to baseline levels. In comparison, no change in Lamin B1 levels was observed.

To further probe the occurrence of DNA damage during mESC differentiation on more *in-vivo* mimetic conditions, we cultured mESCs on mouse embryonic fibroblast (MEF) derived matrices (MEFDMs) which induce mESC differentiation ^61^ (Fig. S 2 Ai, ii). In comparison to mESCs cultured on gelatin-coated glass coverslips in the absence of LIF (serving as controls, CTL), in mESCs cultured on MEFDMs for 24 hrs, increase in γH2AX, loss in Oct3/4 levels and induction of Lamin A/C was observed (Fig. S 2 Bi-iii) when normalized to GAPDH levels (which remained lower for mESCs on MEFDM). While Nanog levels remained unchanged, surprisingly, loss in Lamin B1 was observed. In MEFs which possess high levels of Lamin A/C, no DNA damage was observed. Collectively, our results suggest DNA damage induces loss of pluripotency and Lamin A/C expression.

### Temporal loss of pluripotency is associated with mechanoadaptation mediated differentiation in mESCs

Stimulation by differentiation factors cause pluripotent mESCs to differentiate into either endoderm, ectoderm or mesoderm layers. To probe how mechanoadaptation mediated loss in pluripotency and induction of Lamin A/C expression influences mESC fate, these cells were cultured up to 72 hrs on the PA gels and RNA sequencing (RNAseq) performed at 3, 6, 24 and 72 hr time-points. Principal Component Analysis (PCA) plot shows a clear time dependent clustering of mESCs, with no prominent stiffness dependence (Fig. 4 A). Though DNA damage and Lamin A/C induction was also seen in ETO treated cells, this point did not co-cluster with any of the other time-points. Compared to pluripotent controls (+LIF), temporal RNAseq profiling revealed stemness genes including *Pou5f1* ^62,63^, *Sox2* ^64^, *Nanog* ^65,66^ and *Klf4* ^67^ were maintained till 6 hrs, but were lost completely at the 24 hr time-point (Fig. 4 B). Similar trend was also observed in western blots of Oct3/4 and Nanog (Fig. 4 Ci, ii). Concomitant with loss in pluripotency, several known lineage-specific markers were upregulated at the 72 hr time point. These included endoderm markers such as *Cldn6* ^68,69^, *Gata4* ^70^, *Sox17* ^71^ and *Foxa2* ^72,73^, mesoderm markers such as *Tbx6* ^74,75^, *Twist1*^76–78^ and *Snai1* ^79,80^, and ectoderm markers such as *Nes* ^81^ and *RunX2* ^82^ (Fig. 4 B, Fig. S 3 A). Intriguingly, expression of extraembryonic ectodermal markers such as *Krt8* ^83^, *Wnt7b* ^84^ and *Gata3* ^85^ was also observed. While these results are indicative of heterogeneous differentiation, quantification of the average Z-score values across the three stiffnesses revealed most prominent induction of mesoderm lineage (0.4), and weaker induction of endoderm (0.14) and ectoderm (0.1) lineages, indicating a shift towards mesodermal lineage. Protein level expression of one marker from each lineage – Gata4 for endoderm, Brachyury (T) for mesoderm and NeuN for ectoderm – showed similar trend with RNAseq profiles (Fig. S 4 Ai, ii). In addition, gene ontology analysis revealed lineage specification is mediated by activation of several common transcription factors such as *Zf5*, *E2F*, *SP3*, *Kaiso*, *Pax4*, *Foxn1* (Fig. S 3 B).

**Figure 4:**
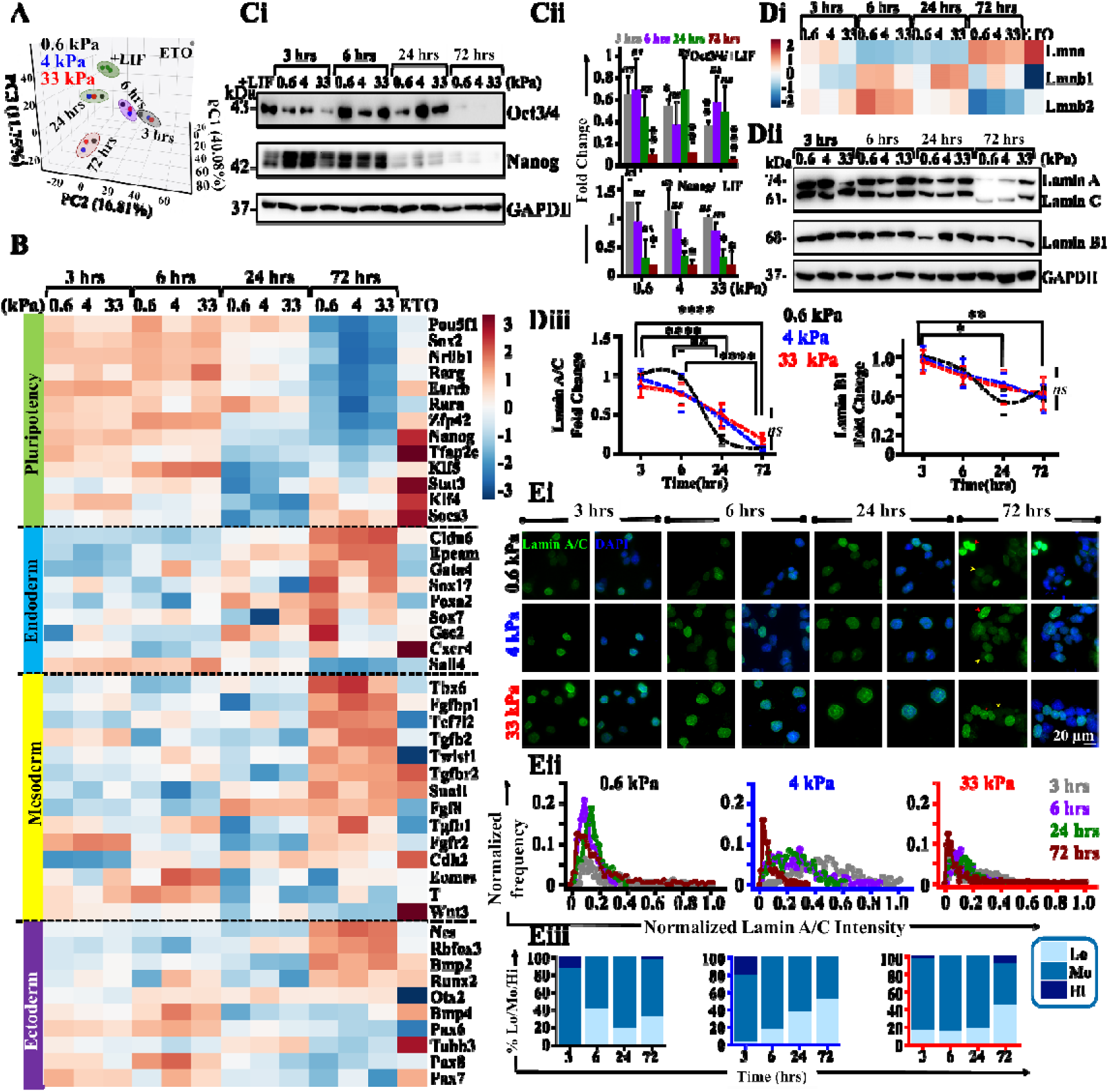
Germ layer differentiation is associated with loss in Lamin A/C: **(A)** 3-D PCA plot representing clusters of mESCs in +LIF condition, Etoposide (ETO) and varying stiffness post 3 hrs, 6 hrs, 24 hrs and 72 hrs of culture based on RNAseq profiles. **(B)** Heatmaps showing temporal evolution of mESC RNAseq profiles of genes involved in pluripotency, endodermal, mesodermal and ectodermal lineage across different stiffnesses and in ETO-treated cells on 0.6 kPa gels. Normalization was carried out with respect to +LIF condition. **(Ci, Cii)** Representative immunoblots and quantification of Oct3/4 and Nanog in mESCs cultured in the presence of LIF and in the absence of LIF on PA gels across different time-points ( independent experiments; error bars represent SEM). Two-way ANOVA with Tukey’s test was used for comparing means with respect to +LIF controls (*-value 0.05, **-value 0.01, ***-value 0.001, ns = non-significant *p*-value 0.05). **(Di)** Heatmaps showing RNAseq profiles of Lamin genes (*LMNA, LMNB1 and LMNB2*) in mESCs cultured on PA gels across different time-points as well as in ETO treated cells on 0.6 kPa gels. Normalization was carried out with respect to +LIF condition. **(Dii, Diii)** Representative immunoblots and quantification of Lamin A/C and Lamin B1 in mESCs cultured on PA gels across different time-points ( independent experiments; error bars represent ±SEM). Quantification of Lamin A/C and Lamin B1 levels were normalized with respect to 0.6 kPa at 3 hrs at different timepoints on the gels. Statistical significance obtained by Two-way ANOVA followed by Tukey’s tests (*p-value <0.05, **p-value <0.01, ****p-value <0.0001, *ns* = non-significant *p*-value >0.05). **(Ei-Eiii)** Representative immunofluorescence images of Lamin A/C co-stained with DAPI in mESCs cultured on PA gels across different time-points showing heterogenous expression of Lamin A/C (red arrowhead: high expression; yellow arrowhead: low expression) (n > 50 cells per condition across *N* = 3 independent experiments). Scale Bar = 20*µm*. Lamin A/C heterogeneity was captured by plotting the normalized Lamin A/C distribution across different time-points. Normalization was done wrt the maximum intensity observed across all conditions and timepoints (n > 50 cells per condition across *N* = 3 independent experiments). Assessment of percentage of low (Lo), moderate (Mo) and high (Hi) Lamin A/C expression across different conditions (see Materials & Methods for details). For all blots, GAPDH served as loading control. See also Figure S 3-S 5.

Temporal RNAseq profiling of Lamin genes across the 4 time-points revealed an increase in *Lmna* gene but drop in *Lmnb1* and *Lmnb2* genes as expected in differentiating stem cells (Fig. 4 Di). Although Lamin A/C is known to enhance differentiation of MSCs ^32^, surprisingly however, protein level analysis shows a gradual decline in both Lamin A/C and Lamin B1 across all the gels, prominently at the 72 hr time-point (Fig. 4 Dii, iii) with considerable heterogeneity in Lamin A/C expression in each condition (Fig. 4 Ei, Red arrowhead: Lamin A/C high; Yellow arrowhead: Lamin A/C low). Lamin A/C intensity distribution revealed an increase in the fraction of Lamin A/C low (Lo) cells at 72 hrs across all the gels (Fig. 4 Eii, iii), with ∼30%, 50% and 40% Lamin A/C low cells at 72 hrs on 0.6 kPa, 4 kPa and 33 kPa gels respectively (Fig. 4 Eiii). However, long term culture of mESCs upto 7 days on the stiff 33 kPa gels, exhibited a recovery of Lamin A/C levels after its initial decline at 72 hrs (3 days) (Fig. S 4B), consistent with previous findings that shows that Lamin A/C loss primes mESCs towards cardiac-specific gene expression ^31,33^. Since, genome integrity is associated with an intact nuclear envelope devoid of blebs and rupture ^42,86^, evaluation of the nuclear lamina using confocal image of Lamin B1, revealed no detectable nuclear blebbing until the 24 hr timepoint on the stiff gels. However, mESCs were seen to exhibit nuclear wrinkling at 72 hrs of culture on these gels (Fig. S 4C).

Since, *LMNA* deficient embryonic hearts have been shown to exhibit cell cycle perturbations during DNA damage ^87^, we next went on to evaluate the cell cycle distribution of mESCs on the PA gels at different time-points. Quantitative analysis showed that sub-G1 phase significantly increased with time – from <20% cells in +LIF mESCs to ∼40% cells at 72 hrs and ∼80% upon ETO treatment (Fig. S 5 Ai, ii). The Sub-G1 cells at 72 hrs may correspond to that of low Lamin A/C expressing mESCs at the same time-point on 33 kPa gels (Fig. 4 Ei-iii), suggesting that transient Lamin A/C loss affects mESC cell cycle distribution on PA gels. Furthermore, quantification of cell survival using Annexin V-PI staining revealed ∼60% live mESCs at 72 hr time-points on all the gels compared to ∼85% live cells in presence of LIF (Fig. S 5 Bi, ii). While temporal RNASeq profiling revealed activation of few apoptosis genes (*Casp8, Casp6, Casp7*) at 72 hrs time-point (Fig. S 5 C), Annexin-PI staining shows no significant increase in apoptotic cells but % necrotic cells were dramatically increased at 72 hr time-point and upon ETO treatment (Fig. S 5 Bi, ii).

Since embryogenesis is mediated by cell-matrix adhesions and mechanical forces ^88^ that control complex mechano-signaling cascades via integrins ^89^, we examined the temporal profile of several genes involved in mechanoadaptation. RNAseq profiles show clear upregulation of several such genes at 72 hrs such as integrin subunits – *ItgaV, Itga3, Itga11, Itgb1, Itgb3, Itgb5, Itgb8*, integrin like kinase (*Ilk*), genes involved in integrin mediated cell adhesion – *Tln1, Vcl, Gsn, Tsn2, Parva, Parvb, Acta1, Acta2, Actg1*, myosin genes – *Mylk, Myl6, Myl9, Myl12a, Myl12b*, along with *Cdh1* and *Cdh2* (Fig. S 4 D). Few LINC complex genes were also upregulated at 72 hrs such as *Sun2, Syne2* and *Lmna* (Fig. S 4 D), thus implicating an integrin dependent mechanoadaptation response that mediates mESC differentiation at 72 hrs. Together, these results suggest that loss in pluripotency in mESCs is associated with transient loss of Lamin A/C protein expression leading to early differentiation (72 hrs) into all three germ layers (with a prominent shift towards mesodermal lineage) through activation of mechano-signaling pathways.

### mESC differentiation is associated with ATR activation

While mESCs have a robust DNA damage response (DDR) mechanism to counter intrinsic and extrinsic assaults to the genome, compared to their differentiated counterparts ^90^, accumulation of DNA damage can cause apoptosis and differentiation to eradicate them from the stem cell pool ^91–93^. We propose that differentiation is triggered via activation of DDR pathways and DNA repair. Consistent with this notion, temporal immunostaining and quantification of γH2AX foci on mESCs displays a steady level upto 24 hrs and then a sharp decline at the 72 hr time-point (Fig. 5 Ai, ii). Similarly, 53BP1 quantification shows a significant increase at 6 hrs which then notably diminished at the 72 hr time-point (Fig. 5 Bi, ii), indicative of a temporal resolution of DNA damage. To further confirm our observations, immunoblotting of mESCs also revealed a significant drop in γH2AX as well as pRPA2 (Single strand break marker) levels at the 72 hr time-point. (Fig. 5 C, Fig. S 6 Ai, ii).

**Figure 5:**
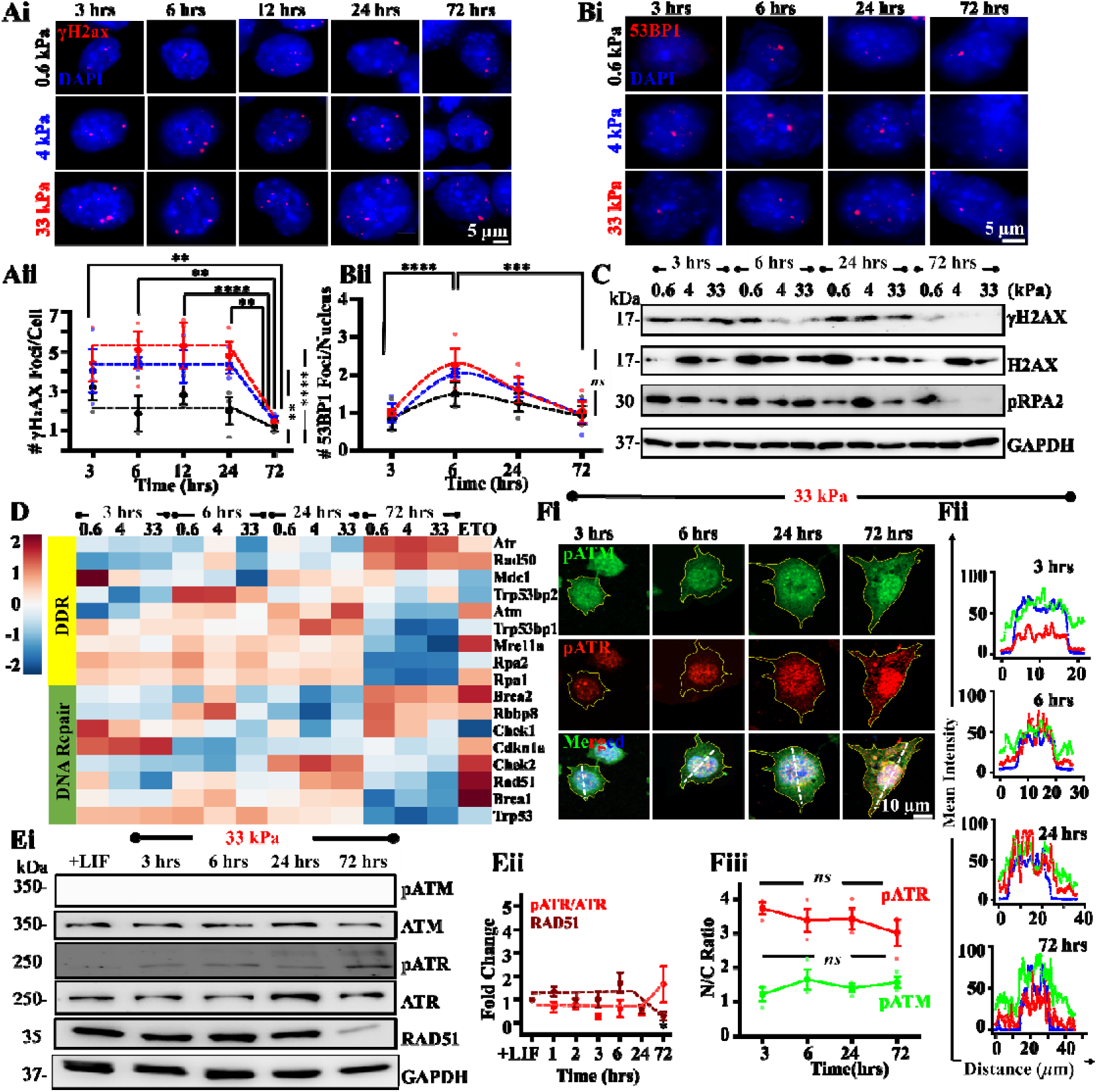
Differentiation involves DNA repair through ATR activation and nuclear enrichment: (Ai, Aii) Representative images of temporal evolution of mESCs immunostained for γH2AX co-stained with DAPI on varying stiffness and quantification of γH2AX foci count per nucleus ( cells per condition across *N* = 3 independent experiments). Scale Bar = 5*µm.* Error bars represent SD. Two-way ANOVA with Tukey’s test was used for comparing means (**-value 0.01, ****-value 0.0001). **(Bi, Bii)** Representative images of temporal evolution of mESCs immunostained for DDR factor, 53BP1, co-stained with DAPI on varying stiffness and quantification of 53BP1 foci count per nucleus ( cells per condition across *N* = 3 independent experiments). Scale Bar = 5*µm.* Error bars represent SD. Two-way ANOVA with Tukey’s test was used for comparing means (*** p-value 0.001, ****-value 0.0001, *ns* = non-significant-value 0.05). **(C)** Immunoblots representing temporal expression of DDR factors, γH2AX/H2AX and pRPA2 of mESCs on varying stiffness. **(D)** Heatmaps showing temporal evolution of mESC RNAseq profiles of genes involved in DNA damage response (DDR) factors and DNA repair across different stiffnesses and in ETO-treated cells on 0.6 kPa gels. Normalization was carried out with respect to +LIF condition. **(Ei, Eii)** Immunoblots representing temporal expression of DDR factors (pATM/ATM, pATR/ATR) and DNA repair factor (RAD 51) in the presence of LIF and in the absence of LIF on 33 kPa PA gels. Quantification of pATR/ATR, ATR, ATM and RAD51 levels were done with respect to +LIF condition at different timepoints across *N* = 3 independent experiments. Error bars represent ±SEM. One-way ANOVA with Tukey’s test was used for comparing means (*p-value ≤0.05). **(Fi, ii)** Representative pATM-Ser1981/pATR-Thr1989/DAPI stained images of mESCs across different timepoints on 33 kPa gels. Scale Bar = 5*µm*. Representative intensity profiles along white dotted lines depict nuclear/cytoplasmic localization of pATM (green) and pATR (red). **(Fiii)** Quantification of nuclear to cytoplasmic (N/C) ratio of pATM (green) and pATR (red) (n > 50 cells per condition across *N* = 3 independent experiments). Error bars represent SD. One-way ANOVA with Tukey’s test was used for comparing means (*ns* = non-significant p-value>0.05). For all blots, GAPDH served as loading control. See also Figure S 6 A-D and Movies S 1-3.

RNAseq profiles of early responders of DNA damage revealed upregulation of *ATR* and *RAD50*, downregulation of *ATM, Mre11a, MDC1, Trp53bp1, RPA1, RPA2, Atrip,* and upregulation of downstream DNA repair genes including *Brca2* and *Rbbp8* at 72 hr time-point (Fig. 5 D). These observations motivated us to investigate the functional roles of ATM and ATR during stiffness mediated mechanoadaptation in regulating lamin levels. No phosphorylated ATM (pATM, Ser1981) bands were detected on 33 kPa gels across the different time-points, notwithstanding the fact that ATM levels remained unchanged across these conditions when compared to +LIF condition (Fig. 5 Ei, Fig. S 6 B). However, pATM was detected in immunostained images of mESCs cultured on 33 kPa gels (Fig. 5 Fi), and exhibited uniform spatial distribution (Fig. 5 Fii, iii). In contrast to ATM, phosphorylated ATR (pATR, Thr1989) was detected from 3 hr onwards (Fig. 5 Ei, ii Fig. S 6 B). Furthermore, quantification of nuclear to cytosolic (N/C) ratio revealed increased nuclear localization of pATR across all time-points (Fig. 5 Fiii) with greater nuclear enrichment on the stiffer 33 kPa gels (Fig. S 6 C). Near complete loss in RAD51 levels at the 72 hr time-point (Fig. 5 D, Ei, ii) may correspond to resolution of DNA damage and mESC differentiation (Fig. 5 A, B and C). Taken together, these results suggest that DNA damage-induced differentiation is associated with ATR mediated DDR activation followed by DNA repair.

Furthermore, since RNAseq analysis of mESCs revealed an upregulation of mesodermal genes, we analyzed the effect of inhibiting DNA repair on development of the heart, a mesodermal organ, which is also mechanically active during its formation. The zebrafish heart forms as a tube and develops into a two-chambered contractile organ by 24 hours post fertilization (hpf). The heart then undergoes morphogenetic movements resulting in a left-ward ventricle and right-ward atrium and chamber ballooning by ∼72 hpf ^94^. Zebrafish embryos exposed to ATM and ATR inhibitors (KU55933 and VE821 respectively) for 24 hours showed mild body curvature defects at 48 hpf (Fig. S 6 Di, ii, n=71 for KU and n=73 embryos for VE) and defects in chamber size and ballooning (Fig. S 6 Dii). Following 24 hours exposure to inhibitors, larvae were allowed to recover in inhibitor-free medium for another 24 hours. However, the defects in heart morphogenesis worsened by 72 hpf, wherein the ventricle and atrium failed to attain their characteristic ballooning and left-right positioning in the larvae (Fig. S 6 Diii, n=35 for KU and n=44 embryos for VE, Supp. movies S1 to S3). Interestingly, in several embryos cardiac contractility was absent at 72 hpf (7/35 for KU and 2/44 for VE). These results suggest that ATM and ATR function is required for normal morphogenesis of the developing zebrafish heart and inhibition of ATM/ATR during cardiac morphogenesis causes inability of embryos to recover from the damage.

### ATR regulates mESC survival and differentiation by temporal modulation of Lamin A/C

Physical properties of the nucleus are dictated by Lamin A/C levels with its phosphorylation a Ser22 leading to its degradation ^32,38^, and inducing nuclear softening^39^. While DNA damage induces Lamin A/C expression at early time-points, Lamin A/C was lost at the 72 hr time-point (Fig. 4 D, E). To probe how DNA damage, DDR response and Lamin A/C degradation are temporally correlated, we tracked γH2AX/H2Ax, pATR/ATR, pCHK1/CHK1, and pLamin A/C (S22)/Lamin A/C over a period of 1 week on 33 kPa gels (Fig. 6 Ai, ii). Similar to ETO treatment (Fig. 3 C), temporal kinetics revealed γH2AX activation at 1 hr precedes Lamin A/C induction at the 3 hr time-point. Lamin A/C induction was associated with CHK1 activation. Subsequent drop in Lamin A/C correlated with increase in pLamin A/C, while pATR levels remained constant. However, increase in pATR after 72 hr was associated with rescue in Lamin A/C levels and concomitant reduction in pLamin A/C levels.

**Figure 6:**
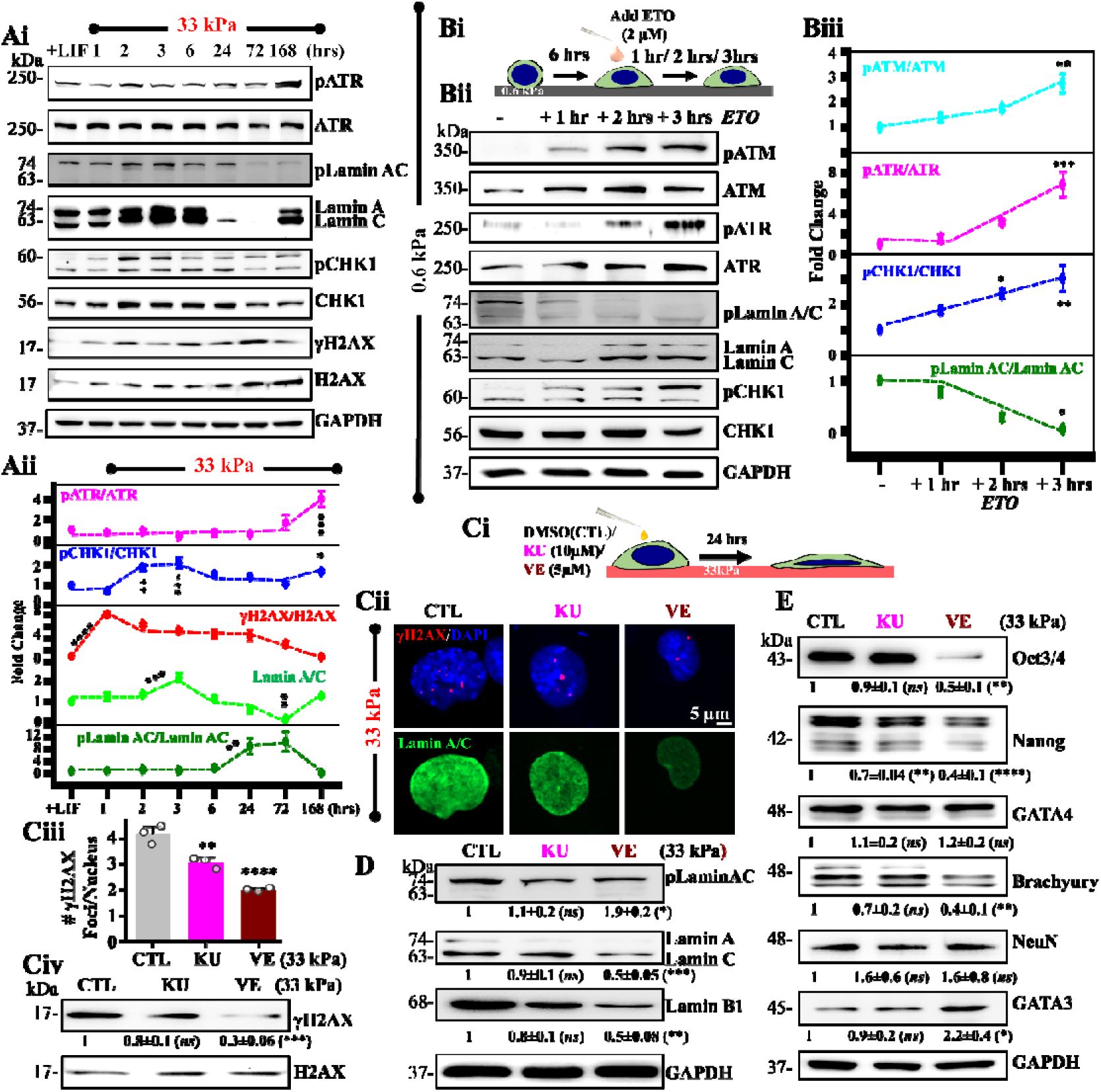
ATR regulates Lamin expression: (Ai, Aii) Immunoblots representing temporal evolution of pATR/ATR, pLamin A/C/Lamin A/C, pCHK1/CHK1, γH2AX/H2AX in the presence of LIF and in the absence of LIF on 33 kPa PA gels upto Day 7 (168 hrs). Quantification of mean ±SEM of pATR/ATR, pCHK1/CHK1, γH2AX/H2AX, Lamin A/C and pLamin A/C/Lamin A/C normalized with respect to +LIF condition ( independent experiments). One-way ANOVA with Tukey’s test was used for comparing means with respect to +LIF controls (*-value 0.05, **-value 0.01, ***-value 0.001, ****-value 0.0001). **(Bi, ii, iii)** Cells were allowed to attach onto 0.6 kPa gels for 6 hrs prior to ETO addition, and then cultured for upto 3 hrs. Representative immunoblots showing temporal evolution of pATM/ATM, pATR/ATR and pCHK1/CHK1, and its quantification of mean SEM ( independent experiments). One-way ANOVA with Tukey’s test was used for comparing means with respect to untreated controls (*-value 0.05, **-value 0.01, ***-value 0.001). **(Ci-iv)** Experimental setup for studying effect of KU-55933 (KU) and VE-821 (VE) on Lamin A/C. Cells were cultured on 33 kPa gels for 24 hrs. Representative immunostained images of γH2AX co-stained with DAPI and Lamin A/C on mESCs cultured in the presence and absence of KU and VE. Bar plots show quantification of γH2AX foci per nucleus in these conditions ( cells per condition across *N* = 3 independent experiments). Scale Bar = 5*µm.* Error bars represent SEM. Unpaired student t-test was used for comparing means between control and treated samples (**p-value <0.01, ****p-value <0.0001). Representative immunoblots showing quantification of mean ±SEM of γH2AX/H2AX (*N* = 3 independent experiments). One-way ANOVA with Tukey’s test was used for comparing means with respect to untreated control (***p-value <0.001, *ns* = non-significant p-value>0.05). **(D)** Representative immunoblots of mESCs on 33 kPa gels in the presence and absence of KU and VE and quantification of mean ±SEM showing pLamin A/C/Lamin A/C, Lamin A/C and Lamin B1 expression (N = 3 independent experiments). One-way ANOVA with Tukey’s test was used for comparing means with respect to untreated controls (\**p*-value <0.05, **p-value <0.01, ***p-value <0.001, *ns* = non-significant). **(E)** Representative immunoblots of mESCs on 33 kPa gels in the presence and absence of KU and VE and quantification of mean ±SEM showing Oct3/4, Nanog, GATA4, Brachyury, NeuN and GATA3 expression (N = 3 independent experiments). One-way ANOVA with Tukey’s test was used for comparing means with respect to untreated control (\**p*-value <0.05, **p-value <0.01, ****p-value <0.0001, ns = non-significant). For all blots, GAPDH served as loading control.

Since both ATM and ATR have been linked to regulation of nuclear envelope integrity and lamin levels ^43,47,95^, we assessed the role of ATM and ATR in regulating Lamin A/C levels. In mESCs cultured in the presence of ETO, though both ATM and ATR were activated, ATR/CHK1 activation was more prominent and correlated with drop in pLamin A/C and increase in Lamin A/C levels (Fig. 6 Bi-iii). To test if ATR regulates Lamin A/C by modulating Lamin A/C phosphorylation, we performed experiments with KU and VE on 33 kPa gels (Fig. 6 Ci). Both inhibitors led to drop in γH2AX foci counts and levels (Fig. 6 Cii-iv). However, only ATR inhibition led to increase in pLamin A/C and decrease in Lamin A/C levels, suggesting that ATR stabilizes Lamin A/C by preventing this phosphorylation (Fig. 6 Cii, D). Even Lamin B1 levels were reduced upon ATR inhibition.

Finally, to investigate the role of ATR activity in mESC cell cycle distribution, viability and differentiation, we analyzed the percentage cell cycle distribution and observed increased SubG1 cells upon VE treatment indicative of a perturbed cell cycle pattern that corresponds to significant reduction in cell viability (Fig. S 7 A, B). Furthermore, assessment of mESC pluripotency and differentiation revealed prominent reduction in Oct3/4 and Nanog levels upon ATR inhibition coupled with significant increase in the extraembryonic ectoderm marker GATA3 and decrease in the mesoderm marker Brachyury (T) (Fig. 6 E). In comparison, GATA4 and NeuN remained unchanged. Taken together, in addition to its role in DDR, our observations suggest that ATR regulates mESC survival and differentiation by modulating Lamin A/C in a temporal manner.

## Discussion

Genome integrity has been associated with transmission of physical forces to the nucleus leading to nuclear envelope deformation, blebbing and rupture in cancer cells ^40,42^. Lamin A/C is a crucial component of the nuclear envelope with Lamin A/C deficiency causing laminopathies ^96^, increased DNA damage ^28,97,98^, and defective mechanotransduction ^30^. Additionally, drug induced DNA damage alters chromatin condensation state resulting in alterations in the biomechanical properties of the nucleus through DDR signaling, thus establishing a link between genome integrity, DNA damage signaling, chromatin condensation and nuclear mechanics ^99^. Here, using FN coated PA gels of varying stiffness, we report stiffer matrices trigger greater DNA damage in mESCs through increased nuclear compression. We show that DNA damage brings about loss in pluripotency and Lamin A/C induction at early time-points, thus functioning as a mediator of early differentiation. Differentiation of mESCs at the 72 hr time-point was associated with early induction of DNA damage response factors (γH2AX, 53BP1 and pRPA2) and downstream DNA repair factors (RAD51). Of the two DNA damage response (DDR) factors ATM and ATR, we establish a role for ATR in modulating and stabilizing Lamin A/C expression by inhibiting its phosphorylation. Collectively, our results demonstrate how stiffness-dependent DNA damage drives differentiation through ATR-dependent Lamin regulation (Fig. 7).

**Figure 7:**
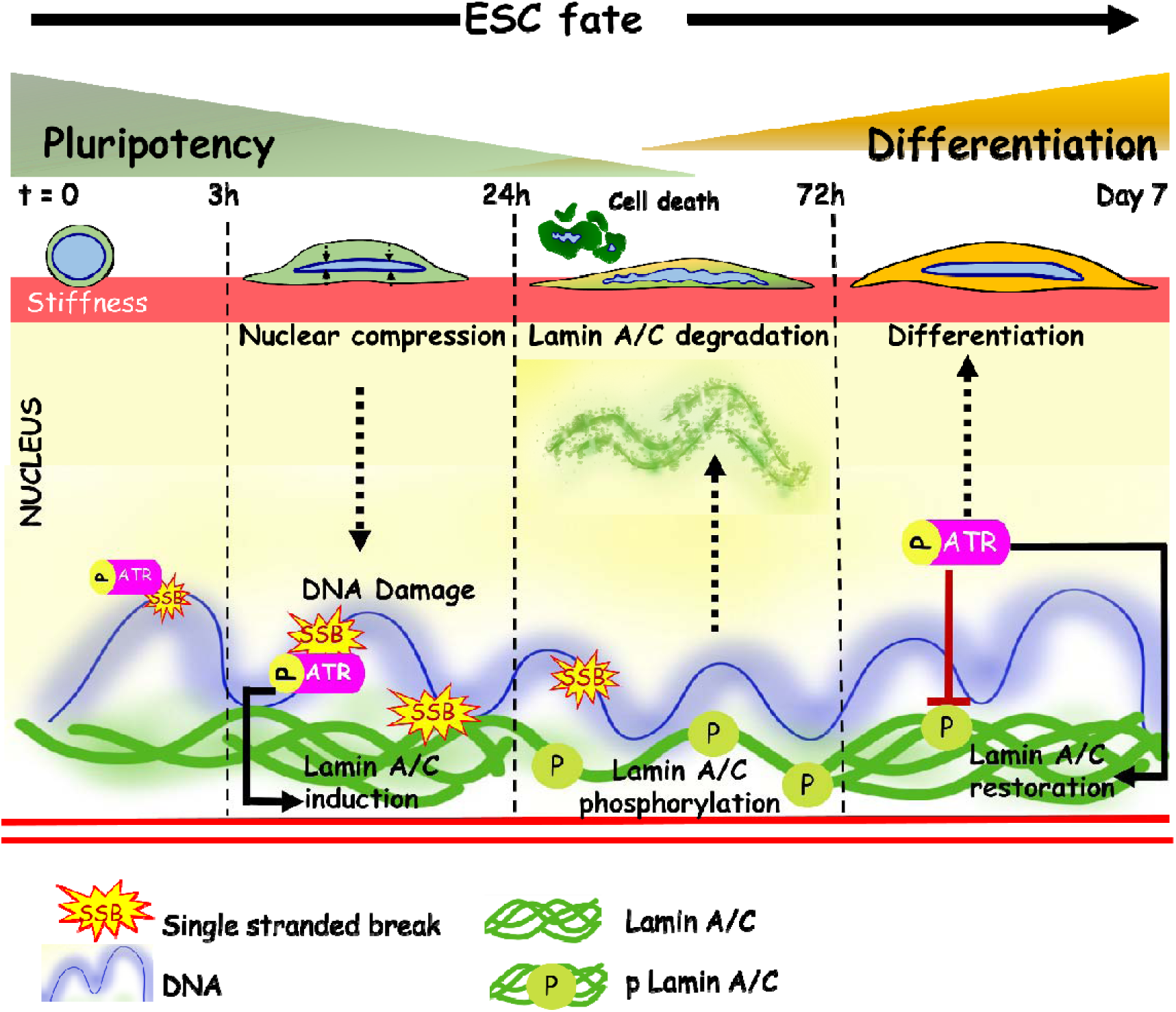
Schematic representation depicting stiffness-dependent differentiation in mouse ESCs. Stiffness-dependent mechanoadaptation induces nuclear compression leading to DNA damage (SSBs), activation of ATR and ATR-mediated lamin A/C modulation. Differentiation involves ATR-mediated lamin A/C stabilization by inhibition of Lamin A/C phosphorylation at early time-points followed by gradual loss of Lamin A/C and cell death. ATR activation in surviving cells at later timepoints restores Lamin A/C.

ATR is activated by single stranded breaks (SSBs) during replication stress that arise due to stalled replication forks and at resected DSBs ^11,100^. As part of the DDR pathway, ATR directly phosphorylates the checkpoint kinase, CHK1 at Ser-345 ^101^. High proliferative capacity of mouse ESCs have been previously shown to contribute to increased accumulation of SSBs and involves ATR mediated activation of H2AX and CHK1 ^53,54^. To assess the nature of DNA damage induced by stiffness, we performed alkaline comet assay which detects both SSBs and DSBs, and neutral comet assay which exclusively detects DSBs. Though both SSBs and DSBs were observed, significantly higher stiffness-dependent increase in olive tail moment of alkaline comets compared to neutral comets suggests that stiffness primarily induces SSBs in mESCs and is correlated with ATR activation. While SSBs are most commonly induced by oxidative stresses^102–104^, insensitivity of γH2AX to the ROS inhibitor NAC indicates that stiffness—induced DNA damage is ROS independent.

Stem cell maintenance and differentiation are regulated by several components of the stem cell niche. Cell-ECM interactions are known to regulate stem cell self-renewability and differentiation^105^. While collagen-I (Col-I) has been reported to maintain mESC self-renewability^106^, fibronectin (FN) has been associated with self-renewability as well as differentiation^107–109^. MEFDMs of different composition, organization and stiffness have been shown to induce distinct mESC germ layer commitment ^61^. In addition to its association with stem cell self-renewability and lineage commitment^19,32,60,110^, matrix stiffness has also been implicated in regulating genomic integrity (ploidy) of mESCs ^48^. Our findings establish a direct link between stiffness mediated mechanoadaptation and DNA damage in mESCs on FN coated PA gels. While MSCs and cancer cells undergo nuclear compression, DNA damage and nuclear envelop rupture during confined migration^40,42,111,112^; we show for the first time that DNA damage in mESCs is attributed to stiffness-dependent nuclear compression and subsequent nuclear wrinkling. Stiffness-dependent differences in nuclear compression observed as early as 3 hrs can be attributed to stiffness-dependent alterations in actomyosin contractility as Blebb treatment on 33 kPa reduced nuclear compression and DNA damage, and MnCl_2_ treatment on 0.6 kPa gels led to increased nuclear compression and DNA damage. Our results are consistent with earlier reports which showed suppression of DNA damage in embryonic hearts by lowering of contractility mediated nuclear deformation^87^.

Literature reports have documented opposite effects of DNA damage in inducing differentiation. While genotoxic DNA damaging agents have been shown to disrupt myogenic differentiation^113^, physiological DNA damage induced by caspase 3 has been shown to promote differentiation^114^. Moreover, confined migration has been reported to cause impaired cell cycle and delayed differentiation in myoblasts^115^. LIF maintains *in-vitro* mESC self-renewability through the STAT3 pathway ^116^, and has been implicated as a mechano-inhibitor ^48^. In mESCs, LIF withdrawal triggers an exit from pluripotency and induction of apoptosis ^117^. However, mESCs on soft substrates maintain pluripotency even in absence of LIF by downregulation of traction forces ^60^. Using RNA sequencing analysis and western blotting we show temporal downregulation of most stemness markers accompanied by upregulation of differentiation genes involved in germ layer commitment, in the absence of LIF. In line with previous observations ^117,118^, temporal cell survivability assay on mESCs show ∼ 40% dead cells at 72 hr time-point. These observations suggests that a proportion of mESCs undergo cell death which could be triggered due to LIF withdrawal for longer duration. Since we also observed time dependent impairment of cell cycle and loss in mESC survivability with ∼60% Sub-G1 cells, the cells undergoing mixed lineage differentiation at the 72 hr time-point likely correspond to the surviving cells where DNA damage in resolved. Consistent with this, expression of the DNA repair protein RAD51 reduced at 72 hrs.

While conflicting reports exist on Lamin A/C expression in mESCs^119,120^, our results reveal non-monotonic alterations in Lamin A/C levels during differentiation. Basal levels of Lamin A/C observed in +LIF conditions is consistent with its role in maintenance of naïve pluripotency and prevention of premature differentiation^33^. By temporally profiling γH2AX and Lamin A/C levels, that stiffness-induced DNA damage precedes induction of Lamin A/C. Induction of Lamin A/C observed at early time-points may contribute to DNA repair in multiple ways. First, stable Lamin A/C localization at the nuclear lamina confers mechano-protection to the genome ^87^. Second, by anchoring damage sites via 53BP1, it can mediate efficient repair^28,121,122^ as evident from ATR/CHK1 activation. Gradual loss of Lamin A/C from 24 hr onwards can be attributed to increased Lamin A/C phosphorylation and degradation^38^ leading to nuclear softening^39^. Given the heterogeneity in Lamin A/C expression, and close correlation between the proportion of low Lamin A/C expressing cells and the proportion of dead cells, it is likely that low Lamin A/C cells may correspond to the genetically defective cells that are eliminated from the stem cell pool during differentiation ^123,124^, with restoration of Lamin A/C at Day 7 corresponding to differentiated cells. Thus, Lamin A/C plays a temporally differential role during stiffness induced mESC differentiation with induction at early timepoints protecting the genome, and loss at intermediate timepoints initiating differentiation.

Apart from the canonical DNA damage response exhibited by ATM and ATR, these factors have been extensively linked to nuclear envelope integrity, regulation of lamins and mechanosensitivity during cancer migration and invasion ^43–47,95^. While ATM has been claimed to be activated by cell stretching^46^, ATR is activated by nuclear compression during interstitial migration^43^. Another study has shown the role of ATR in Lamin A/C phosphorylation at Ser 282 in response to DNA damage, leading to alterations in nucleo-cytoskeletal and chromatin organization ^95^. Here, we show that pATM is uniformly present in nucleus and cytosol, but pATR selectively localizes to the nucleus on stiff gels. Temporal analysis of our study suggests that although pATR levels remain unchanged till 72 hr timepoint, constitutive nuclear localization and pCHK1 activation synchronous with Lamin A/C levels is indicative of a connection between ATR activation and Lamin A/C modulation. Since ATR stabilizes Lamin A/C by inhibiting its phosphorylation, we posit that differential role of Lamin A/C during differentiation is regulated by ATR/CHK1.

ATR is known to be developmentally essential^14^ and has been stated to mediate fate decisions in ESCs during replication stress^125^. Our experiments in zebrafish embryos also show that ATM and ATR function is required for normal morphogenesis of the mechanically contractile zebrafish heart during critical stages of cardiac development. The changes in shape and size of the heart together with the heart beat defects indicate that cell fate specification and cell differentiation processes may be affected. Interestingly, substantial increase in pATR levels on Day 7, in line with Lamin A/C restoration is indicative of its role in initiation of stiffness dependent mESC differentiation. Although inhibition of ATR activity does not significantly change; levels of GATA4, Brachyury and NeuN, increase in levels of GATA3, the extraembryonic ectoderm was observed. This points towards the idea that the ATR/CHK activity could be responsible for maintaining mesodermal lineage specification during stiffness induced mechanoadaptation, while its suppression drives extraembryonic ectodermal lineage specification as seen from our RNA seq data and western blots at 72 hr time point. Taken together, our results indicate a novel role of matrix stiffness induced nuclear compression that triggers DNA damage in activating ATR within the nucleus, and driving ESC differentiation through modulation of Lamin A/C.

In conclusion, our results illustrate the role of matrix stiffness in mediating nuclear compression in mESCs leading to DNA damage. Our results also show that such DNA damage not only drives early differentiation and Lamin A/C induction but also activates and localizes ATR to the nucleus, indicative of the role of ATR in modulating lamin expression in mESCs during mechanoadaptation.

## Acknowledgements

Authors acknowledge financial support from the Department of Biotechnology (Govt. of India) (Grant # BT/PR34522/MED/31/417/2019) and intramural funds provided by IIT Bombay. TR was supported by a CSIR fellowship (Grant # 09/087(0927)/2017-EMR-I) (Govt. of India). Authors would also like to thank IRCC, IIT Bombay for providing Bio-AFM and confocal microscopy facilities, and DST FIST sponsored FACS facility. Authors are deeply grateful to Prof. Anirban Banerjee and Prof. Santanu Kumar Ghosh, Department of Biosciences and Bioengineering, IIT Bombay for their critical inputs and suggestions.

## Author contributions

Conceptualization: T.R., S.K., S.N., C.S.P., S.S.; Methodology: T.R., S.G., N.P., S.D., N.T., C.W.K., S.S.P., P.S., S.G., L.K.S., D.T.S., S.N., C.S.P., S.K, S.S.; Software: T.R., S.D., N.T., C.W.K., P.S., D.T.S., C.S.P., S.N., S.K., S.S.; Validation: S.S.; Formal analysis: T.R., S.D., S.G., N.T., C.S.P., S.K., S.S.; Investigation: T.R., S.K., S.N., S.S.; Resources: S.S.; Data curation: T.R., S.G., N.P., S.D., N.T.; S.S.P.; C.S.P.; S.K., S.S.; Writing - original draft: T.R., S.S.; Writing - review & editing: T.R., N.P., S.D., C.W.K., N.T., C.S.P., S.N, S.K., S.S.; Visualization: T.R., S.S.; Supervision: S.S.; Project administration: S.S.; Funding acquisition: S.S.

