## Supplementary Figures and legends for "Nuclear compression-mediated DNA damage drives ATR-dependent Lamin expression and mouse ESC differentiation"

### Supplemental Information titles and legends

### Figure S1

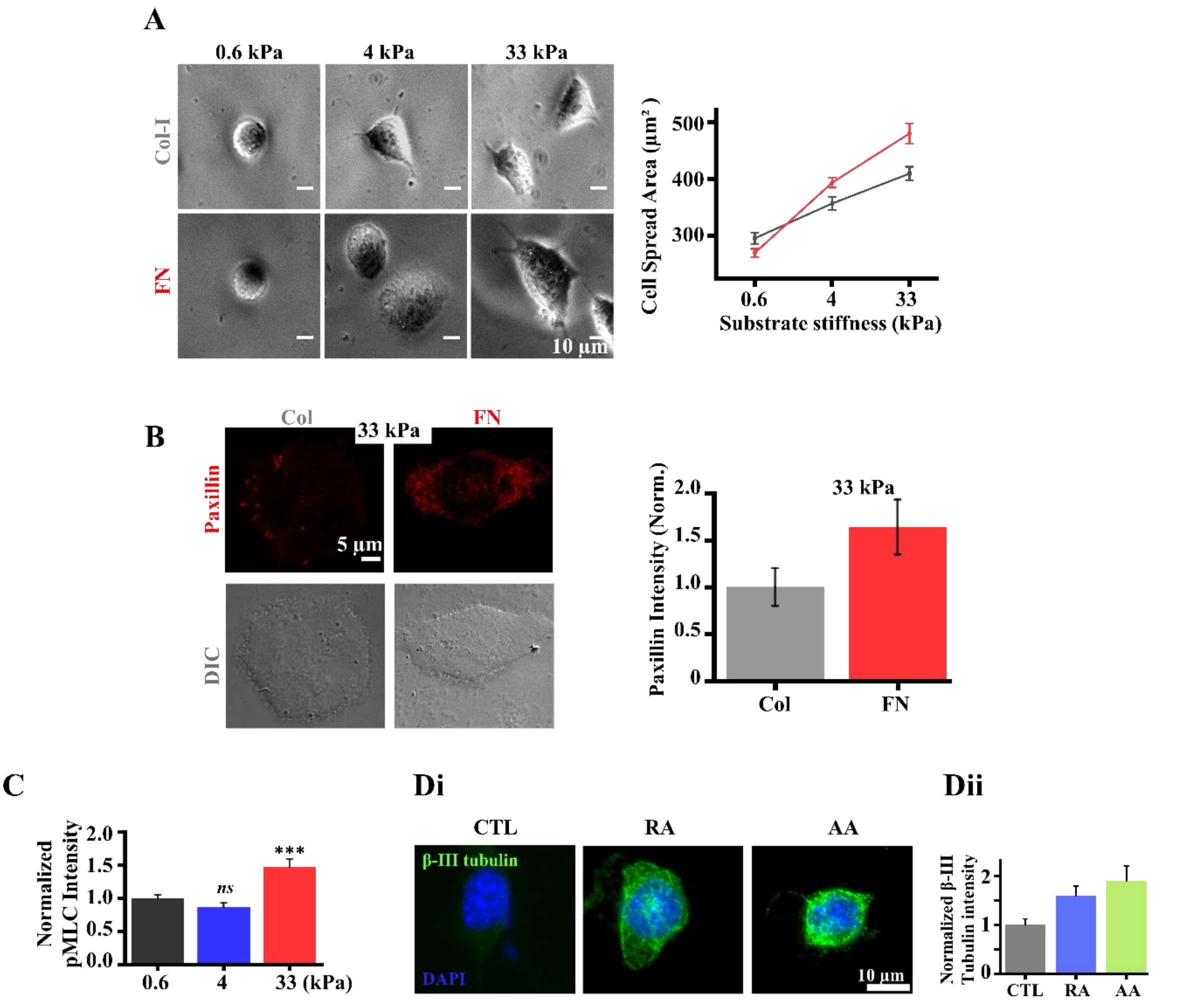


**Supplementary Figure S1: mESCs mechanoadapt on Fibronectin (FN) but not on Collagen-I (Col-I) coated PA gels: (A)** Representative phase contrast images of mESCs on PA gels of varying stiffness coated with either Col-I or FN and quantification of cell spreading ($n \geq 30$ cells per condition across $N = 3$ independent experiments). Scale Bar = $10\mu m$. Error bars represent SEM. **(B)** Representative paxillin stained and DIC images of mESCs on Col-I vs FN coated 33 kPa gels and quantification of paxillin intensity normalized with Col-I intensity ($n \geq 30$ cells per condition across $N = 3$ independent experiments). Scale Bar = $5\mu m$. Error bars represent SEM. **(C)** Quantification of pMLC intensities normalized with respect to 0.6 kPa condition ($n \geq 30$ cells per condition across $N =$ 3 independent experiments). Error bars represent normalized SEM. Statistical significance obtained by unpaired t-test (***$p$-value $\leq$0.001, *ns* = non-significant $p$- value >0.05). **(Di, ii)** β-III tubulin/DAPI immunostained images of mESCs on 0.6 kPa at 24 hr time-point in presence and absence of RA/AA. Quantification of mean intensities of β-III tubulin normalized wrt untreated condition, CTL. ($n \geq 40$ cells per condition across $N = 1$ independent experiment). Scale Bar = 10$\mu$m. Error bars represent normalized SEM.

**Figure S2:**

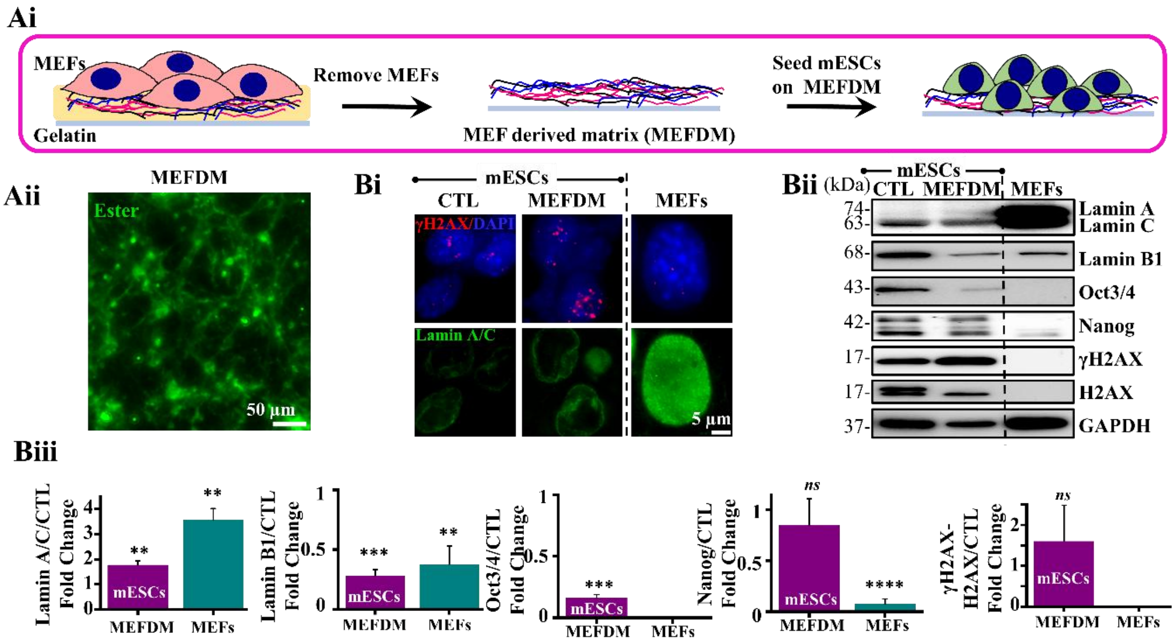


**Supplementary Figure S2: DNA damage induces Lamin A/C in MEFDMs: (Ai, ii)** Schematic representation of MEF derived matrices (MEFDMs) preparation from MEFs and mESC seeding. Representative immunofluorescence image of MEFDM stained with NHS-ester (Scale Bar = 50$\mu m$). **(Bi)** Representative immunofluorescence images of Lamin A/C and γH2AX of mESCs on gelatin coated glass coverslips and on MEFDMs compared to differentiated MEFs. Scale Bar = 5$\mu m$. **(Bii)** Representative immunoblots of Lamin A/C, Lamin B1, Oct3/4, Nanog, γH2AX and H2AX of whole cell mESC lysates isolated from gelatin coated dishes (CTL) and from MEFDM compared to MEF whole cell lysates. **(Biii)** Quantification of Lamin A/C, Lamin B1, Oct3/4, Nanog and γH2AX/H2AX immunoblots of mESCs on either gelatin coated dishes (CTL) or on MEFDMs and of MEFs, normalized with respect to mESCs on gelatin coated dishes across $N$ = 3 independent experiments. Error bars represent SEM. Statistical significance for each condition assessed by Student's t-test compared with CTL (mESCs on gelatin coated dishes) (**$p$-value $\leq$0.01, ***$p$-value $\leq$0.001, ****$p$-value $\leq$0.0001, *ns* = non-significant $p$-value >0.05). For all blots, GAPDH served as loading control.

### Figure S3

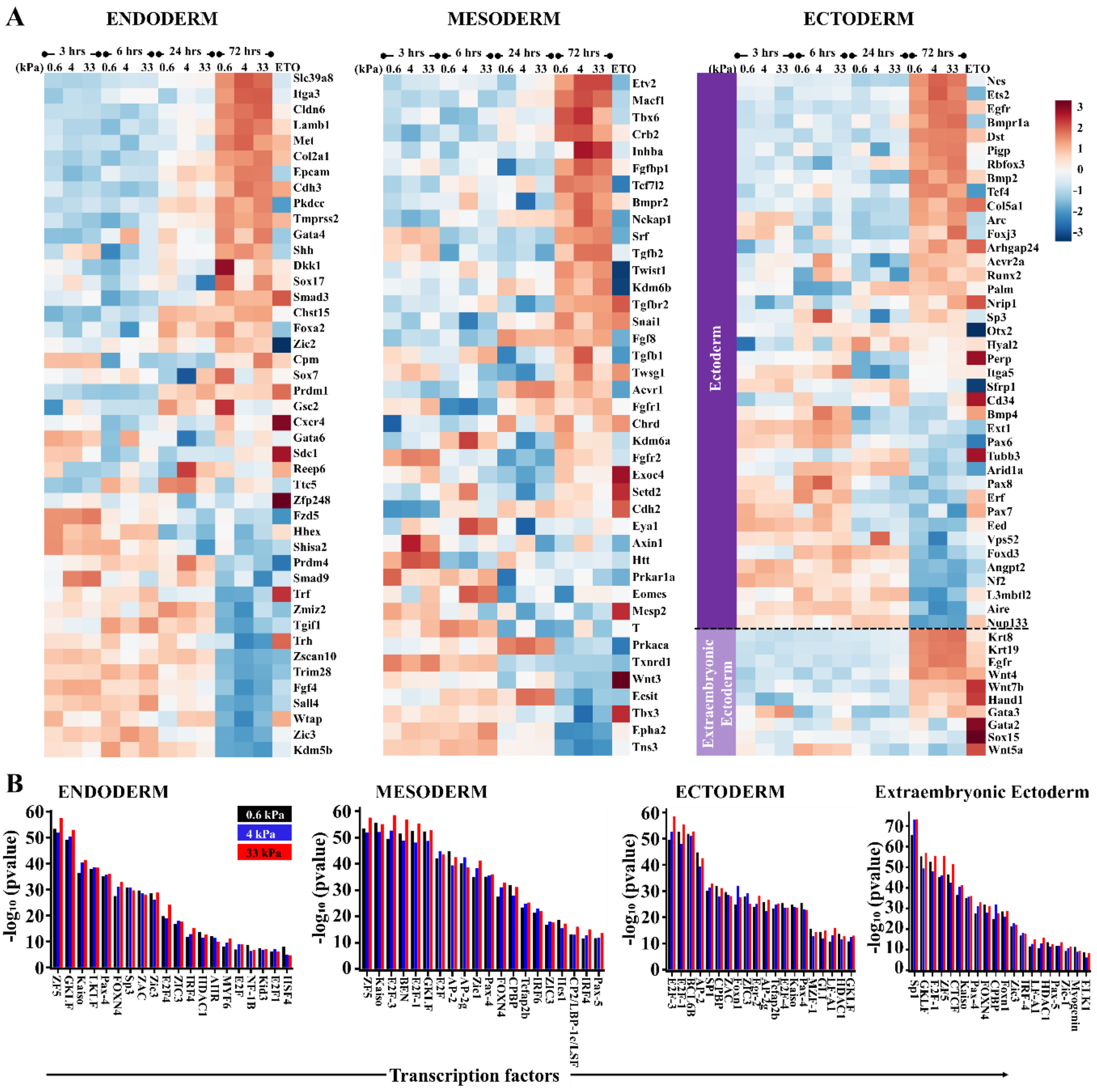


**Supplementary Figure S3: Lineage specification of mESCs during mechanoadaptation: (A)** Heatmaps representing RNAseq profiles of genes associated with endoderm, mesoderm, ectoderm and extraembryonic ectoderm across different stiffnesses and timepoints and on ETO treated mESCs on 0.6 kPa gels. Normalization was carried out with respect to +LIF condition and genes arranged according to descending order of average Z-score values at the 72 hr time-point. **(B)** Top 20 transcription factors ranked according to p-values which are upregulated across the three stiffnesses at the 72 hr time-point represented based on frequency of occurrence during endoderm, mesoderm, ectoderm and extraembryonic ectoderm lineage specification respectively.

### Figure S4

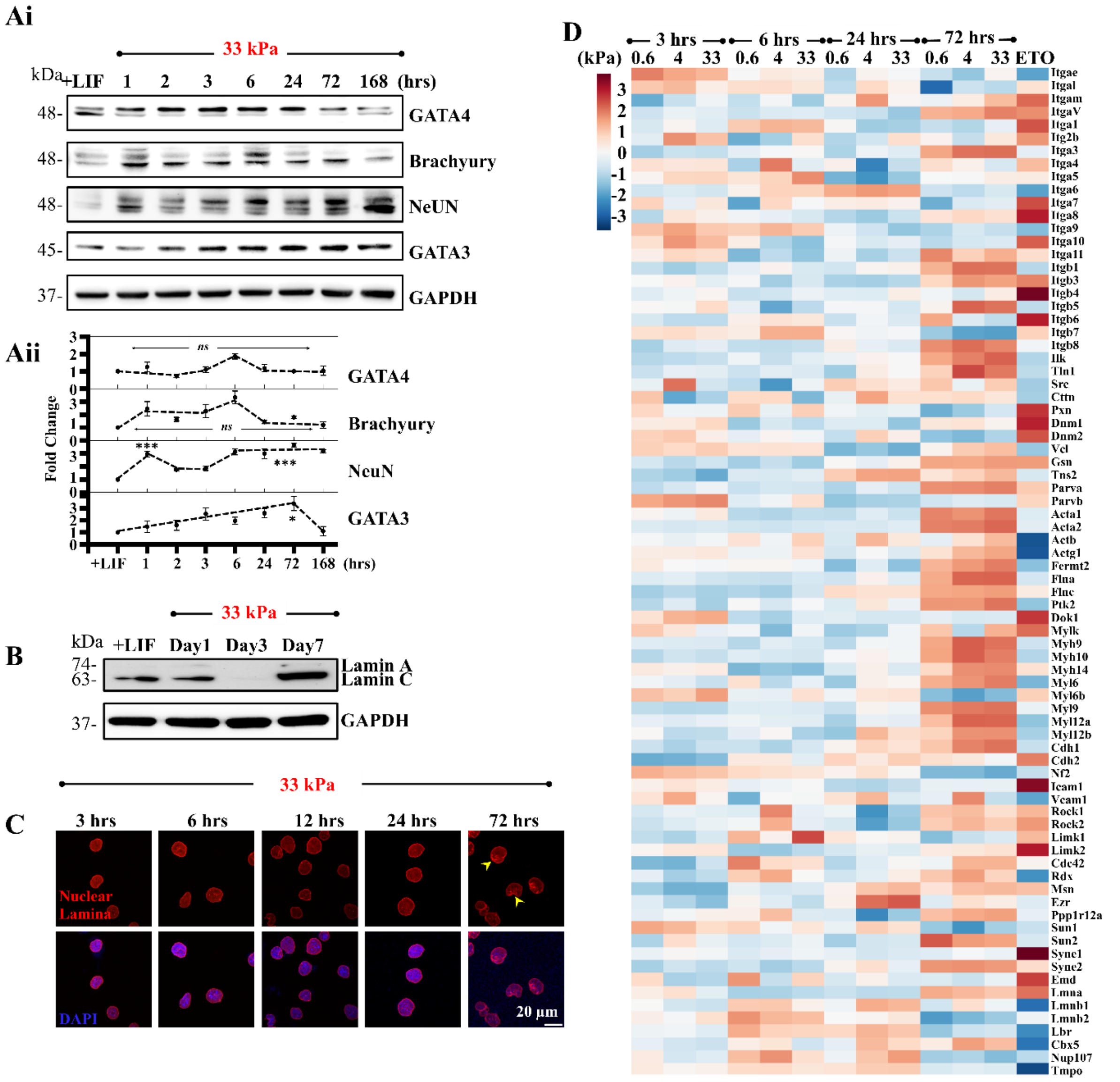


**Supplementary Figure S4: Temporal evolution of mESC pluripotency and Lamins and their association with mechanoresponsive genes: (Ai)** Representative immunoblots showing temporal evolution of endo-, meso-, ecto- and extraembryonic ectodermal markers of mESCs on 33 kPa PA gels. **(Aii)** Quantification of GATA4, Brachyury, NeuN and GATA3 levels with respect to +LIF condition at different timepoints on 33 kPa gels across *N* = 3 independent experiments. Error bars represent ±SEM (**p*-value ≤0.05, ****p*-value ≤0.001, *ns* = non-significant *p*-value >0.05). One-way ANOVA followed by Tukey's test was used for comparing means. (**B**) Representative immunoblot showing Lamin A/C levels on +LIF condition, day1, day 3 and day7 of mESC culture clearly depicting its loss at day 3 and restoration on day7 (*N* = 2 independent experiments). **(C)** Representative immunostained images of Lamin B1 (nuclear lamina) co-stained with DAPI, showing intact nuclei upto 24 hrs and yellow arrowheads showing nuclear envelope wrinkling at 72 hrs **(D)** Heatmap representing RNAseq profiles of genes associated with cell-matrix adhesion, focal adhesion, cell-cell adhesion, migration and nuclear mechanics across different stiffnesses and timepoints and on ETO treated mESCs on 0.6 kPa gels. Normalization was carried out with respect to +LIF condition. GAPDH served as loading control for all immunoblots.

**Figure S5**

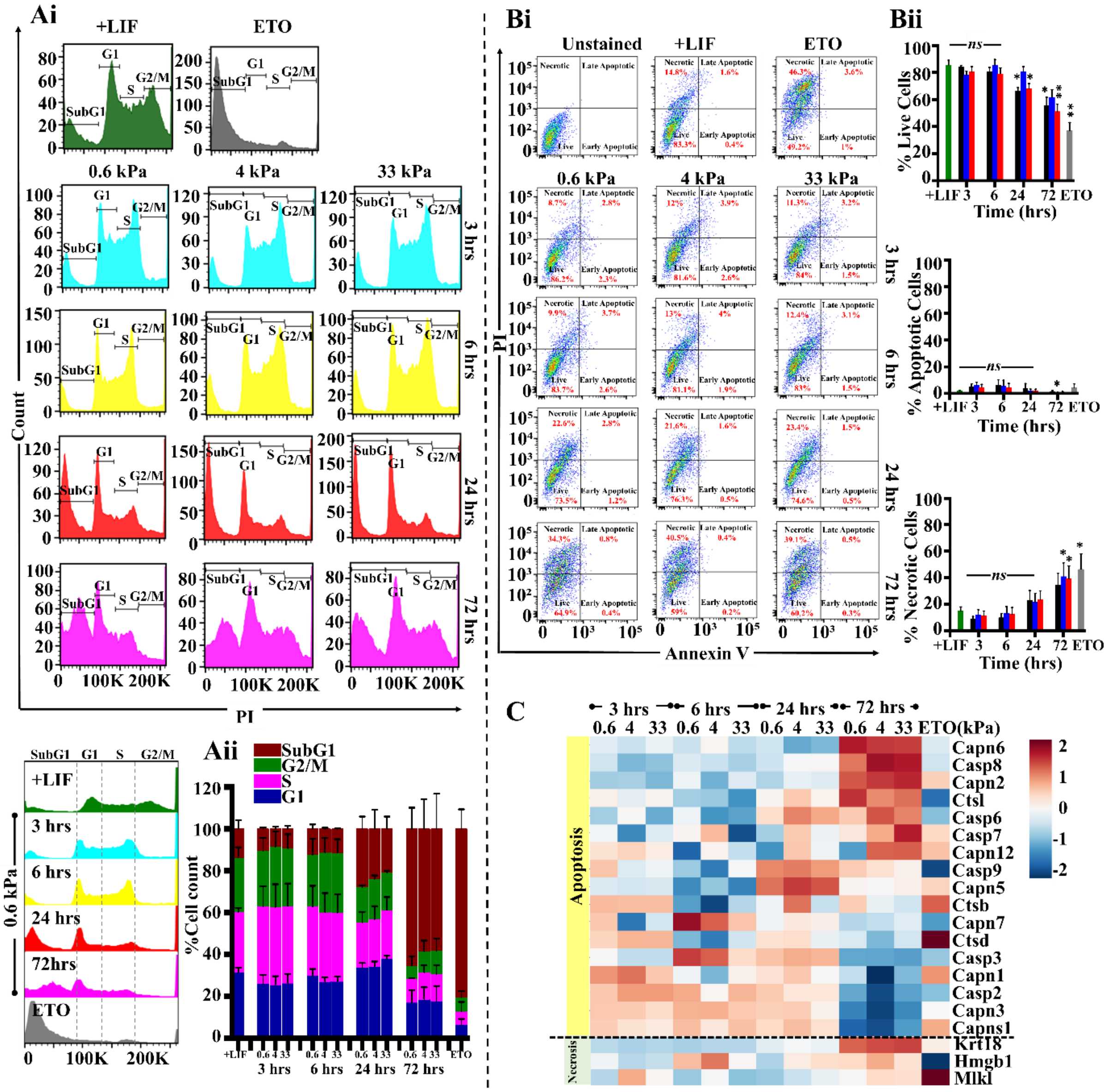

**Supplementary Figure S5: Lamin A/C loss is associated with aberrant cell cycling and increased necrosis: (Ai)** Representative histograms showing flow cytometry analysis of cell cycle distribution of mESCs in +LIF condition, ETO and different timepoints on 0.6 kPa, 4 kPa and 33 kPa gels. Gating was normalized with respect to +LIF condition. **(Aii)** Percentage cell cycle distribution in mESCs in +LIF condition, ETO and PA gels across different time-points (*N* =3 independent experiments). Error bars represent SEM. **(Bi)** Representative plots showing distribution of cell viability by Annexin V/PI staining of mESCs in +LIF condition, ETO and different timepoints on 0.6 kPa, 4 kPa and 33 kPa gels. Gating was done with respect to Annexin-ve and PI-ve mESCs (Unstained). **(Bii)** Bar graphs depicting temporal evolution of percentage live, apoptotic and necrotic mESCs across different stiffnesses and in +LIF condition and ETO-treated cells (*N* = 3 independent experiments). Error bars represent SEM. Statistical significance assessed by Student's t-test compared to +LIF control (**p*-value ≤0.05, ***p*-value ≤0.01, *ns* = anon-significant *p*-value >0.05). **(C)** Heatmaps showing temporal evolution of mESC RNAseq profiles of genes involved in apoptosis and necrosis pathways across different stiffnesses and in ETO-treated cells on 0.6 kPa gels. Normalization was carried out with respect to +LIF condition.

**Figure S6**

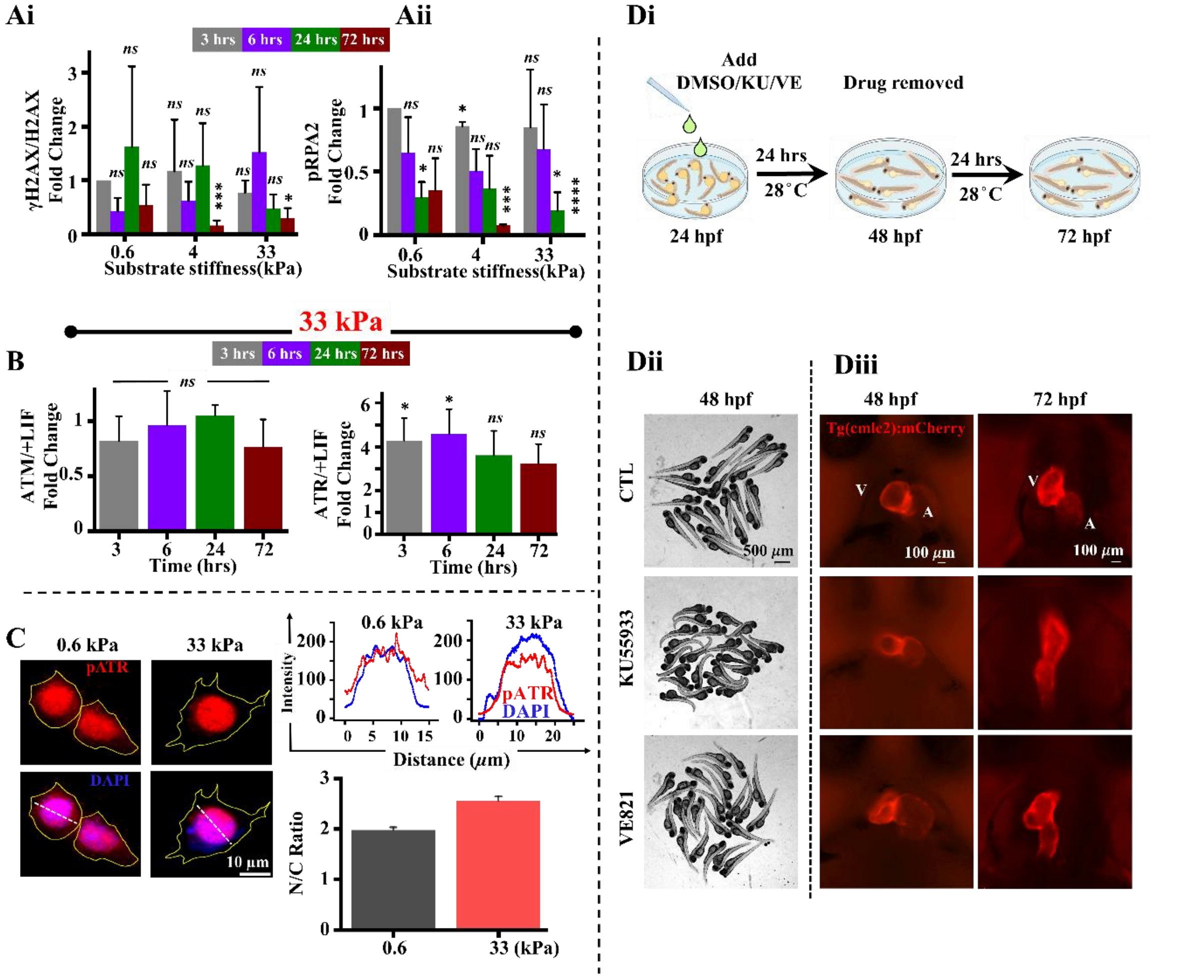


**Supplementary Figure S6: mESC differentiation and zebra fish development and its correlation with DDR factors – ATM and ATR in regulating Lamin levels: (Ai, ii)** Quantification of γH2AX/H2AX and pRPA2/GAPDH levels normalized with respect to 0.6 kPa at 3 hrs at different timepoints on the gels across $N = 3$ independent experiments. Error bars represent normalized SEM. Statistical significance assessed by unpaired Student's t-tests (*$p$-value $\leq 0.05$, ***$p$-value $\leq 0.001$, ****$p$-value $\leq 0.0001$, *ns* = non-significant $p$-value $>0.05$). **(B)** Quantification of ATR and ATM levels with respect to +LIF condition at different timepoints across $N = 3$ independent experiments. Error bars represent normalized SEM. Statistical significance obtained by unpaired Student's t-test (*$p$-value $\leq 0.05$, *ns* = non-significant $p$-value $>0.05$). **(C)** Representative pATR-Thr1989/DAPI stained images of mESCs on 0.6 kPa and 33 kPa gels at the 24 hr time-point. Scale Bar = 10$\mu m$**.** Representative intensity profiles along white dotted lines depict nuclear/cytoplasmic localization of pATR (red) and DAPI (blue). Quantification of nuclear to cytoplasmic (N/C) ratio of pATR (red) ($n \geq 30$ cells per condition across $N = 1$ independent experiment). Error bars represent SEM. **(Di)** Schematic representation of zebra fish embryo experimental setup. **(Dii)** Phase contrast images of zebra fish embryos before (24 hpf) and after ATMi/ATRi treatments (KU55933 and VE-821 respectively). Scale Bar = 500$\mu m$. **(Diii)** Fluorescence images of *tg(cmlc2):m-Cherry* tagged zebra fish heart at 48 hpf and 72 hpf before and after KU and VE treatments depicting developmental defects in heart in comparison to DMSO control (CTL) Scale Bar = 100 $\mu m$. See also supplemental movies S1-S3.

### Figure S7

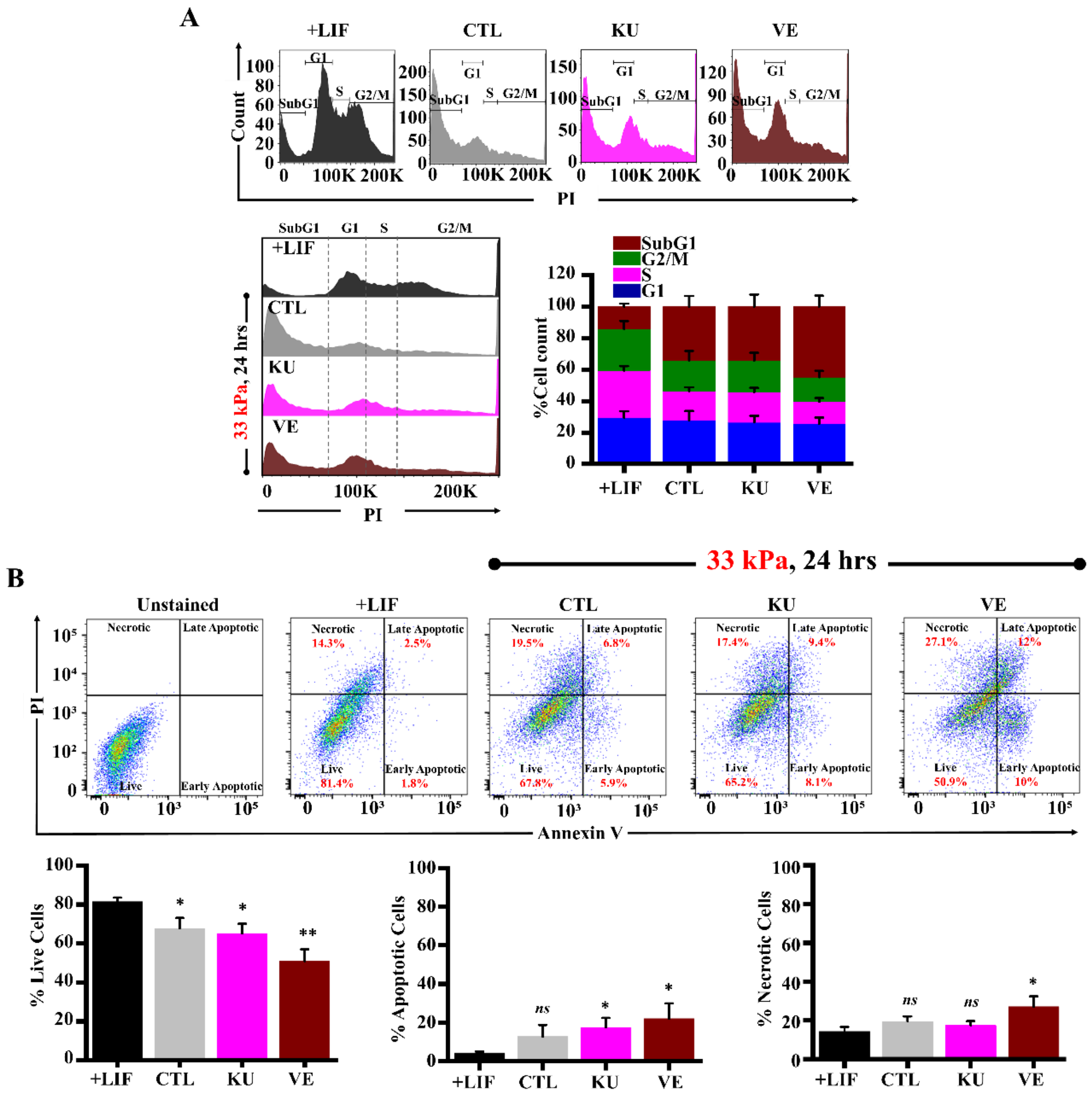


**Supplementary Figure S7: ATR phosphorylation regulates Lamin A/C levels: (A)** Representative histograms showing flow cytometry analysis of cell cycle distribution of mESCs in +LIF, untreated control and in presence of either ATM or ATR inhibitors (KU or VE respectively) on 33 kPa gels. Percentage cell cycle distribution in mESCs in +LIF, CTL, KU and VE on 33 kPa PA gels (*N* =3 independent experiments). Error bars represent SEM. **(B)** Representative plots showing distribution of cell viability by Annexin V/PI staining of mESCs in +LIF condition, CTL, KU and VE on 33 kPa gels. Gating was done with respect to Annexin -ve and PI -ve mESCs (Unstained). Bar graphs depicting percentage live, apoptotic and necrotic mESCs in +LIF, CTL, KU and VE conditions (*N* = 3 independent experiments). Error bars represent SEM. Statistical significance assessed by Student's t-test compared to +LIF control (**p*-value ≤0.05, ***p*-value ≤0.01, *ns* = non-significant *p*-value >0.05).
